# Random birth order and bursty Notch ligand expression drive the stochastic AC/VU cell fate decision in *C. elegans*

**DOI:** 10.1101/588418

**Authors:** Simone Kienle, Nicola Gritti, Jason R. Kroll, Ana Kriselj, Yvonne Goos, Jeroen S. van Zon

## Abstract

Cells in developing organisms must robustly assume the correct fate in order to fulfill their specific function. At the same time, cells are strongly affected by molecular fluctuations, i.e. ‘noise’, leading to inherent variability in individual cells. During development, some cells are thought to exploit such molecular noise to drive stochastic cell fate decisions, with cells randomly picking one cell fate out of several possible ones. Yet, how molecular noise drives such decisions is an open question. We address this question using a novel quantitative approach to study one of the best-understood stochastic cell fate decisions: the AC/VU decision in *C. elegans* gonad development. Here, two initially equivalent cells, Z1.ppp and Z4.aaa, interact, so that one cell becomes the anchor cell (AC) and the other a ventral uterine precursor cell (VU). It is thought that the symmetry is broken when small molecular fluctuations are amplified into cell fate by positive feedback loops in the Notch signaling pathway. To identify the noise sources that drive the AC/VU decision, we used a novel time-lapse microscopy approach to follow expression dynamics in live animals and single molecule FISH to quantify gene expression with single mRNA resolution. We found not only that random Z1.ppp/Z4.aaa birth order biased the decision outcome, with the first-born cell typically assuming VU fate, but that the strength of this bias and the speed of the decision decreased as the two cells were born closer together in time. Moreover, we find that the Notch ligand *lag-2/Delta* exhibited strongly stochastic expression already in the two mother cells, Z1.pp/Z4.aa. Combining experiments with mathematical models, we showed that the resulting asymmetry in *lag-2/Delta* levels inherited by the daughter cells, Z1.ppp/Z4.aaa, stochastic symmetry breaking when both cells are born at similar times. Together, our results suggest that two independent noise sources, birth order and stochastic *lag-2/Delta* expression, are exploited to amplify noise into cell fate in a manner that ensures a robust decision.

## Introduction

Perhaps the most striking feature of development is the emergence of highly complex cellular structures that are nevertheless strikingly reproducible from embryo to embryo. Yet, paradoxically, many cell fate decisions that underlie development are known to be inherently stochastic [1]. For example, the first cell fate decision in the mouse embryo, between trophectoderm and primitive endoderm fate, is stochastic[2]. Moreover, stochastic cell fate decisions are integral to generating the wide range of terminally differentiated cells in development of the nervous system[3–8] and epithelial tissues [9].

It is commonly assumed that stochastic cell fate decisions are driven by small fluctuations on the molecular level, referred to as ‘noise’[10] that are amplified into one cell fate or another by feedback loops in the underlying gene regulatory network[1,11]. This picture is largely based on studies performed on single-celled organisms that in response to changing environmental conditions assume different cellular states in a stochastic manner[12,13]. However, due to the difficulty of studying such stochastic, dynamical processes in multi-cellular organisms, the mechanisms underlying stochastic cell fate decisions in development remain poorly characterized

Moreover, stochastic cell fate decisions likely face different constraints in developing organisms than stochastic transitions in cellular states in single-celled organisms. An important consideration in developing organisms is that cells generated by stochastic cell fate decisions are often needed for subsequent development and hence have to be generated within limited time window, typically on the timescale of hours. It is an open question how a fundamentally stochastic process can guarantee an outcome within such a time window. In addition, the noise sources that drive stochastic cell fate decisions in development are not well-characterized. In single-celled organisms, noise in gene expression has emerged as a key driver[14]. However, in multi-cellular systems noise could be generated in sources, such as cell-cell communication, that have no equivalent in single-celled organisms. Finally, it remains an open question how strong the variability in these noise sources is. In particular, it is not known whether the initial fluctuations have to be sufficiently strong in order to be amplified into cell fate within the time window allotted by the developmental program.

Here, we address these questions in the AC/VU decision, a classical stochastic cell fate decision that occurs during development of the reproductive system in the nematode *C. elegans*. During the late-L1 larval stage, two distantly related precursor cells, Z1.pp and Z4.aa, divide in the gonad to produce four cells (Fig. 1a). The two outer cells, Z1.ppa and Z4.aap, assume Ventral Uterine (VU) fate and contribute to the uterus. However, the two inner cells, Z1.ppp and Z4.aaa, assume either Anchor Cell (AC) or VU fate in a stochastic and mutually exclusive fashion[15], with the AC subsequently inducing vulva and uterine development[16]. Genetic analysis has revealed that Z1.ppp and Z4.aaa communicate via Notch signaling through the Notch receptor *lin-12/Notch* and Notch ligand *lag-2/Delta*[17–19]. These studies suggested that Notch signaling functions as a feedback loop in two ways (Fig. 1b). First, activation of LIN-12/Notch receptor in one cell by binding of LAG-2/Delta ligands from the other cell leads to inhibition of *lag-2/Delta* gene expression in the first cell[18,19]. This occurs through Notch-mediated degradation of the transcription factor, HLH-2, required for *lag-2/Delta* expression[20,21]. Second, activated Notch signaling is thought to induce further *lin-12* expression [20]. As a result of this feedback loop, small random differences between the two cells, e.g. in LIN-12/Notch or LAG-2/Delta level, are thought to be amplified in a stochastic manner into a state where one cell only expresses *lag-2/Delta* (AC fate), while in the other *lag-2/Delta* expression is fully inhibited by Notch signaling (VU fate). This mechanism of stochastic symmetry breaking by mutually exclusive Notch inhibition is widely conserved and responsible for spatial pattern generation, e.g. during bristle patterning in Drosophila[22,23].

**Figure 1.**
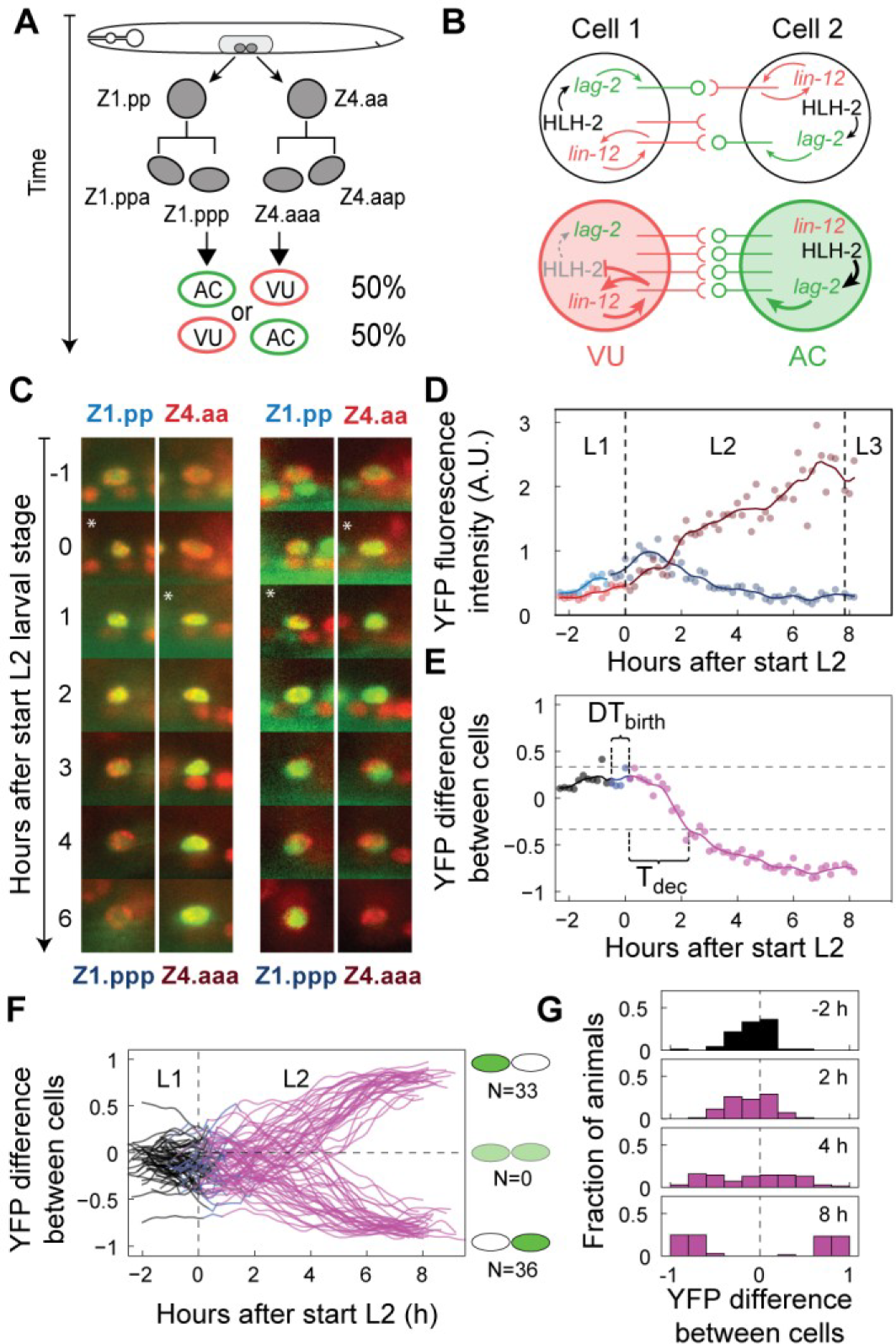
Single-animal expression dynamics during the AC/VU decision. **(a)** Stochastic specification of anchor cell (AC) and ventral uterine (VU) fate in the Z1.pp(p)/Z4.aa(a) lineage. **(b)** Model of AC/VU fate specification by signaling through the Notch receptor LIN-12/Notch (red) and ligand LAG-2/Delta (green). The transcription factor HLH-2 induces *lag-2* expression and is degraded upon activation of LIN-12/Notch. Small, random initial variability is amplified into cell fate by feedback loops in the Notch signaling network. **(c)** *lag-2* expression dynamics in an animal where either Z1.ppp (left column) or Z4.aaa (right column) assumes AC fate. Activation of *lag-2* expression is measured using a *lag-2p::nls::yfp* reporter (green) that localizes to the nucleus. Cells are identified using a body-wide nuclear marker (red). A star indicates the first time point after cell division. **(d)** *lag-2p::nls::yfp* fluorescence level in Z1.pp/Z1.ppp (light/dark red) and Z4.aa/Z4.aaa (light/dark blue) cells as a function of time in a single animal. Markers are individual measurements and lines sliding averages. Dashed lines are time of ecdysis, indicating larval stage transitions. **(e)** Normalized fluorescence difference (*YFP*_z1.pp(p)_-*YFP*_z4.aa(a)_)/(*YFP*_z1.pp(p)_ + *YFP*_z4.aa(a)_) for the animal in (d). Δ*T*_birth_ is the time between birth of Z1.ppp and Z4.aaa. The time to decision, *T*_dec_, is the time between the birth of the second-born cell and the moment YFP fluorescence intensity in one cell is two times that in the other cell (corresponding to the grey dotted lines). Color gives the stage of the AC/VU decision: both mother cells Z1.pp/Z4.aa present (black), one daughter cell, Z1.ppp or Z4.aaa, born (cyan) and both daughter cells born (magenta). **(f)** Overview of *lag-2p::nls::yfp* dynamics for a wild-type population of *n* = 69 animals. Shown is normalized difference. **(g)** Histogram of normalized fluorescence difference at different times. Over time, the distribution transforms from unimodal to bimodal, with high *lag-2p::nls::yfp* expression restricted a single cell.

Even though the AC/VU decision is well-studied genetically, the stochastic expression dynamics of the key regulators *lin-12/Notch* and *lag-2/Delta* has not been studied in single animals. As a consequence, it remains an open question how efficiently Notch signaling breaks the symmetry between initially identical cells. Given that the AC and VU cells are indispensable for further development, their fates need to be assigned in time. Yet, it is not known how much time is required on average for the AC/VU decision to be completed, i.e. when an AC emerges that fully represses *lag-2* expression in its neighbor, nor how variable this decision time is from one animal to another. In order to ensure a sufficiently rapid stochastic decision, it is not only important how rapidly initial variability is amplified, but also how strong this initial variability is. Observations in a limited number of animals revealed that the order in which the Z1.ppp and Z4.aaa cells are born is random, with the first-born cell often assuming AC fate[20], implicating birth-order as a noise source driving the decision. However, how strongly birth order biases cell fate outcome and whether other noise source operate in parallel is currently not known.

A main hurdle in addressing these questions is that they require quantifying stochastic dynamics during the AC/VU decision. Here, we overcome these challenges using two complementary approaches. First, we use single molecule FISH (smFISH) to quantify expression of *lin-12/Notch* and *lag-2/Delta* with single mRNA precision in fixed animals[24,25]. However, stochastic decisions are strongly history-dependent and need to be followed in time in single animals. Here, we used a new time-lapse microscopy approach we developed recently[26] to directly observe expression dynamics of transcriptional *lag-2/Delta* reporter during AC/VU decision. Using this approach, we found that AC/VU was driven by changes in *lag-2/Delta*, but not lin-*12/Notch* expression dynamics. Surprisingly, we found strong and stochastic expression of *lag-2/Delta* already in the Z1.pp and Z4.aa, the mothers of Z1.ppp and Z4.aaa. Combining smFISH and time-lapse experiments with mathematical modeling, our results suggested that bursty, stochastic *lag-2/Delta* expression in Z1.pp and Z4.aa acts as a parallel noise source to drive the AC/VU decision in their daughters, Z1.ppp and Z4.aaa, when they are born at similar times and, hence, cannot efficiently use the birth order as their primary noise source.

## Results

### Stochastic single-cell expression dynamics during the AC/VU decision

Expression patterns of the Notch receptor *lin-12* and its ligand *lag-2* in Z1.pp(p) and Z4.aa(a) cells have been characterized during the AC/VU decision only at single time point in a small number of animals[19,27]. To understand how initial variability in these cells is amplified into cell fate it is essential to follow these processes directly in time in single animals. For this, we used a novel time-lapse microscopy approach that we recently developed to visualize single-cell dynamics in moving and feeding *C. elegans* larvae during their entire ~40hr development[28]. To measure *lag-2* expression dynamics, we employed a *lag-2p::2xNLS::yfp* transcriptional reporter (*arIs131[lag-2p::2XNLS::YFP]*) that expresses nuclearly localized YFP in cells, such as P6.p, the AC and the distal tip cells, known to express *lag-2[29]*. We combined this reporter with a transgene, *itIs37[pie-1p::mCherry::H2B::pie-1]*, that expresses nuclearly localized H2B::mCherry in all somatic cells, to identify the Z1.pp(p) and Z4.aa(a) cells in the gonad. In this way, we were able to observe *lag-2p::yfp* dynamics consistent with that expected during the AC/VU decision, with similar YFP levels in newly born Z1.ppp/Z4.aaa cells that became subsequently restricted to either the Z1.ppp or Z4.aaa cell (Fig. 1b).

To confirm that the observed *lag-2p::yfp* dynamics reflected the expression dynamics of the endogenous *lag-2* gene, we characterized *lag-2* expression in fixed, wild-type animals using smFISH as an independent measure (Fig. 2). Overall, in Z1.ppp and Z4.aaa cells we found a similar progression from symmetrical *lag-2* expression that became progressively restricted to one of the two cells (Fig. 2a-c). Moreover, when we examined *lag-2* mRNA levels and YFP fluorescence in *lag-2p::yfp* animals (Supplementary Fig. 1a), we found that most animals (56/65) that exhibited YFP fluorescence also expressed *lag-2*. The small number of animals that showed *lag-2* expression, but no YFP fluorescence (7/65) or the reverse (2/65) are consistent with the longer production time and longer lifetime, respectively, of YFP proteins compared to mRNA. Using smFISH, we found that *yfp* and *lag-2* mRNA levels were positively correlated, with levels of *yfp* mRNA only 2-3 times higher than of *lag-2* (Supplementary Fig. 1b), suggesting that *lag-2p::yfp* was a relatively low-copy insertion. Overall, these observations showed that *lag-2p::yfp* is a faithful reporter of the underlying *lag-2* expression dynamics.

**Figure 2.**
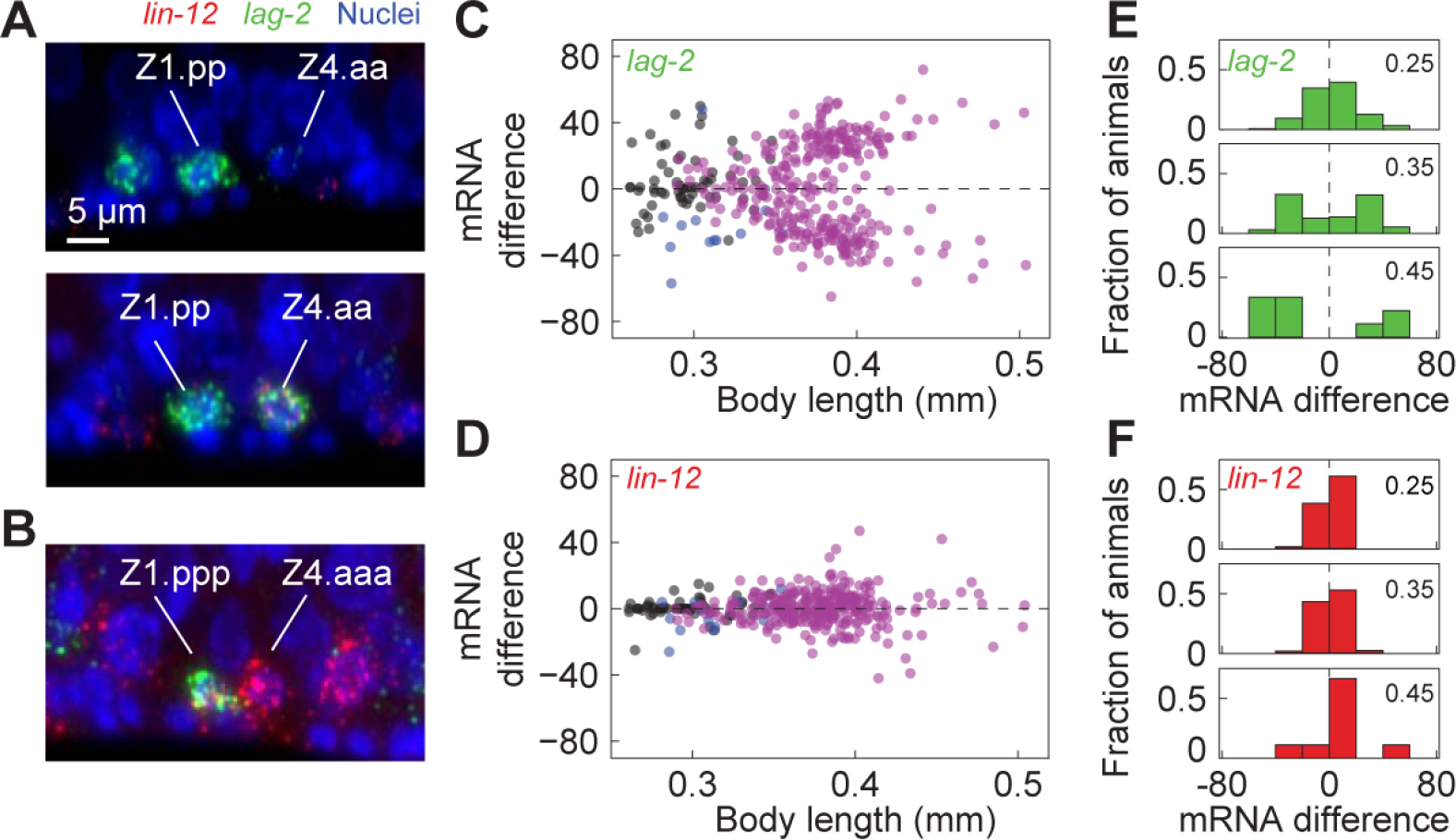
Single-molecule quantification of *lag-2/Delta* and *lin-12/Notch* expression. **(a)**,**(b)** smFISH staining of *lag-2* (green) and *lin-12* (red) mRNA molecules in (a) Z1.pp/Z4.aa mother cells and (c) Z1.ppp/Z4.aaa daughter cells. In animals during the early L2 larval stage *lag-2* expression is highly variable, with animals showing both very different (a, upper panel) or highly similar (a, bottom panel) *lag-2* levels between the mother cells Z1.pp and Z4.aa. In animals at the late L2 stage *lag-2* expression is restricted to either the daughter cell Z1.ppp (b) or Z4.aaa. **(c, d)** Difference in mRNA level, *M*_Z1.pp(p)_ - *M*_Z4.aa(a)_, for (c) *lag-2* and (d) *lin-12* mRNA level as a function of body length. Difference is in units of mRNA molecules. Each marker corresponds to a single animal, with color indicating both mother cells Z1.pp/Z4.aa present (black), one daughter cell, Z1.ppp or Z4.aaa, born (cyan) and both daughter cells born (magenta). Body length is used a measure of developmental stage, with the range observed here corresponding to late L1 to late L2 larval stage. Variability is higher for *lag-2* than *lin-12* expression. In contrast to *lag-2*, no restriction of *lin-12* expression to a single cell is observed. **(f)** Histogram of difference in *lag-2* (green) and *lin-12* (red) mRNA levels at body lengths of 0.25, 0.35 and 0.45mm, calculated for 0.1mm window size.

### Changes in *lag-2*/Delta but not *lin-12*/Notch expression drive the AC/VU decision

Stochastic symmetry breaking between two or more cells through Notch signaling is commonly thought to require feedback mechanisms that result in reciprocal changes in Notch receptor and ligand level, with inhibition of Notch ligand expression accompanied by increased Notch receptor expression[30]. For the AC/VU decision, this feedback is thought to result in *lin-12* Notch receptor levels that are the reverse of *lag-2* Notch ligand levels; low in the AC and high in the VU cell [19]. However, when we examined LIN-12 protein dynamics during the AC/VU decision using a low-copy integrated translation fusion, *wgIs72* [31], we found that LIN-12::GFP levels remained high in both cells, even at the late-L2 stage where we saw clear restriction of *lag-2p::nls::yfp* expression to Z1.ppp or Z4.aaa (Supplementary Fig. 2). Moreover, we found that while *lin-12* expression in wild-type animals showed considerable variability from animal to animal, we found a significant fraction of animals that showed equal *lin-12* mRNA levels at the late-L2 stage where *lag-2* expression was only found in a mutually exclusive manner (Fig. 2b-e). These results suggested that during the AC/VU decision, *lin-12* expression dynamics plays no role in the feedback mechanism, but rather that *lin-12* functions as a passive communication channel to allow feedback through changing *lag-2* ligand levels. For this reason, in our subsequent analysis we focused exclusively on *lag-2* dynamics.

### Single-animal variability in *lag-2*/Delta dynamics

To characterize the animal-to-animal variability in *lag-2* expression dynamics during the AC/VU decision, we quantified the average nuclear *lag-2p::nls::yfp* fluorescence intensities *I*_*Z1*_ and *I*_*Z4*_ in the mother cells Z1.pp/Z4.aa and daughter cells Z1.ppp/Z4.aaa in time in single live animals (Fig. 1d). In all animals, *lag-2p::nls::yfp* fluorescence increased over time until 1-3hr into the L2 larval stage both Z1.ppp/Z4.aaa were born. Subsequently, fluorescence kept increasing in the future AC but decayed in the future VU. To facilitate comparing dynamics between animals, we calculated the fluorescence intensity difference Δ*I* = (*I*_*Z1*_ − *I*_*Z4*_)/(*I*_*Z1*_ + *I*_*Z4*_) between the two cells, with Δ*I* = 0 corresponding to equal expression in Z1.pp(p) and Z4.aa(a) and Δ*I* = ± 1 corresponding to expression fully restricted to the Z1.pp(p) cell or Z4.aa(a) cell, respectively (Fig. 1e). In addition, we used the nuclear H2B::mCherry marker to detect Z1.pp/Z4.aa cell divisions, allowing us to determine birth order as well as time between birth Δ*T*_*birth*_ (Fig. 1e). Finally, we estimated the time for the AC/VU decision to complete, *T*_*decision*_, as the time required for one cell, either Z1.ppp or Z4.aaa, to express the *lag-2p::nls::yfp* reporter at a level twice that of the other, i.e. 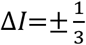, measured with respect to the time of birth of the second-born cell.

Overall, we found that *lag-2p::nls::yfp* dynamics was highly variable between animals. We found extensive variability in dynamics between animals (Fig. 1f,g). In Z1.pp/Z4.aa mother cells and newly-born Z1.ppp/Z4.aaa daughters (~2hr after the start of the L2 larval stage) the intensity difference followed a broad unimodal distribution, indicating continuous variability in reporter level between the two cells. At ~8hr after the start of L2, this initial variability was amplified into a striking bimodal distribution, with two peaks of equal width and height, corresponding to AC fate assumed by Z1.ppp (Δ*I* ≈ 1) or Z4.aaa (Δ*I* ≈ −1). At this stage, no animals were observed with equal reporter expression, (Δ*I* ≈ 0). At intermediate times (~4hr after the start of L2), we observed a very broad, but unimodal distribution of relative reporter levels between Z1.ppp and Z4.aaa. As discussed in more detail below, this largely reflects variability in the decision time *T*_dec_ between animals. In a significant fraction of animals, we observed that during the AC/VU decision the difference in *lag-2p::nls::yfp* expression between the two cells was reversed, i.e. the cell that ultimately assumed AC fate did not initially have the highest fluorescence level (Fig. 1d,e, 40/69 animals). Such reversals only happened once, if at all, in a 3hr period after the birth of the second cell. Finally, we observed very similar distributions when examining the difference in *lag-2* mRNA levels between Z1.ppp and Z4.aaa, as obtained by smFISH (Fig. 2c,e), a further indication that *lag-2p::nls::yfp* reporter dynamics reproduces the expression dynamics of the endogenous *lag-2* gene.

### Birth order biases cell fate outcome for sufficiently large time between births

Experiments where the Z1.pp/Z4.aa lineages were followed in live animals using Nomarski microscopy showed that the first-born cell assumed VU fate in most animals [Karp2006]. Here, we made use of the larger population size in our experiments, as well as our ability to visualize *lag-2* expression dynamics, to examine the role of birth order as a noise source for the AC/VU decision in more detail. We found that Z1.ppp and Z4.aaa were born around the start of the L2 larval stage, with time of birth correlated but still showing substantial variability between the two cells (Fig. 3a). We found that Z1.ppp and Z4.aaa were the first-born cell with equal probability, with Z1.ppp born first in 32/69 animals. We found that the time between Z1.ppp and Z4.aaa births, Δ*T*_birth_, followed an approximately exponential distribution, with Z1.ppp/Z4.aaa born with 30 minutes in most animals but extremes observed up to 1-2hrs (Fig. 3b). Using the *lag-2p::nls::yfp* reporter, we connected birth order to cell fate outcome. Indeed, we found that the first-born cell more frequently assumed VU fate, with the bias towards VU fate increasing with time between birth Δ*T*_birth_ (Fig. 3b). However, for Δ*T*_birth_ <30 min, we observed a significant fraction (13/40 animals) where the first-born cell instead assumed AC fate. Using the *lag-2p::nls::yfp* reporter, we could now also examine how *lag-2* expression dynamics depended on Δ*T*_birth_, the time between Z1.ppp/Z4.aaa birth (Fig. 3c). In general, we found that the time to decision *T*_dec_ showed strong animal-to-animal variability, irrespective of the time between birth. However, on average the time to decision was longer when Z1.ppp and Z4.aaa were born more closely together in time, with the longest decision times *T*_dec_ ≈ 9hr observed only in animals where Δ*T*_birth_ ≤ 20 min. Overall, these results confirm that variability birth order is a source of noise that drives the first-born cell to VU fate, but only with a strong birth order bias and rapid decision time when Z1.ppp and Z4.aaa are born sufficiently far apart in time. This raises the question what other noise sources drive the AC/VU decision when the birth order cue is absent or weak, i.e. Δ*T*_birth_ ≈ 0?

**Figure 3.**
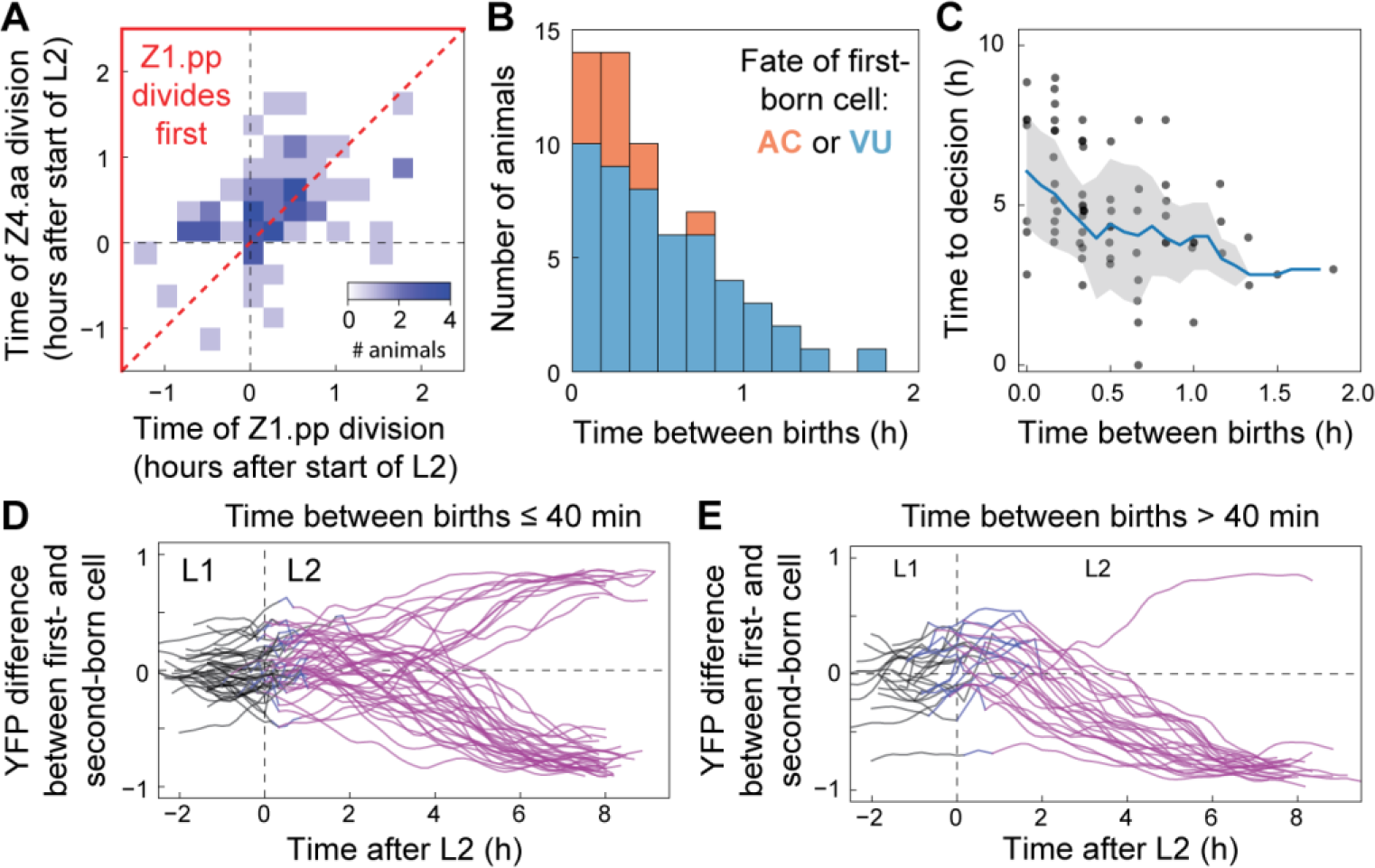
Birth order biases first-born cell to VU fate. **(a)** Histogram of time of Z1.pp and Z4.aa division in single animals. Color indicates the number of animals and the region outlined in red corresponds to animals where Z1.pp divides before Z4.aa. Variability in the time of Z1.pp and Z4.aa division results in variability in birth order and time between Z1.ppp/Z4.aaa births. **(b)** Number of animals where first-born cell assumes VU (blue) or AC (orange) fate as a function of the time between birth of the first- and second-born cell. Birth order bias is strongest for animals where Z1.ppp and Z4.aaa are born far apart in time. **(c)** Time to decision (as defined in Fig. 1e) as a function of time between births. Markers correspond to individual animals. The blue line is the average for all animals with the same time between births, with the grey area indicating the standard deviation. On average, animals where Z1.ppp and Z4.aaa are born close together in time require more time to restrict *lag-2p::yfp:nls* expression to a single cell. **(d,e)** Dynamics of *lag-2p::yfp::nls* in single animals, for animals where Z1.ppp/Z4.aaa are born (d) ≤40 min or (e) >40min apart. Shown is (*YFP*_1_-*YFP*_2_)/(*YFP*_1_ + *YFP*_2_), where *YFP*_1,2_ is the YFP level in the lineage with the first- and second-born, cell respectively. Color indicates stage of the decision: two mother cells (black), one daughter cell born (cyan) and both daughter cells born (magenta).

### Notch-independent, bimodal *lag-2* expression in Z1.pp/Z4.aa mother cells

Gene expression often shows strong, stochastic variability [11,13,32] and hence could function as a noise source for stochastic cell fate decisions. When we examined *lag-2* expression by smFISH, we observed that *lag-2* was expressed already at high levels, ~40 molecules, in the mother cells Z1.pp and Z4.aa, prior to the presumed start of the AC/VU decision in their daughters Z1.ppp and Z4.aaa. Moreover, *lag-2* expression patterns in Z1.pp/Z4.aa were highly variable, meaning that we observed animals with either high or low expression in both Z1.pp and Z4.aa cells as often as animals with high *lag-2* expression in one but almost no expression in the other cell (Fig. 2a,c). Surprisingly, when we measured the distribution of *lag-2* expression levels pooled for both Z1.pp and Z4.aa mother cells, we found a striking bimodal distribution (Fig. 4c). Such bimodal distributions in mRNA level are a hallmark of ‘bursty’ gene expression[32], where cells stochastically switch between periods of high and low expression. In contrast, weaker variability and unimodal distributions were observed both for *lag-2* expression in the daughter cells Z1.ppp/Z4.aaa and for *lin-12* expression at all stages (Supplementary Fig. 3).

**Figure 4.**
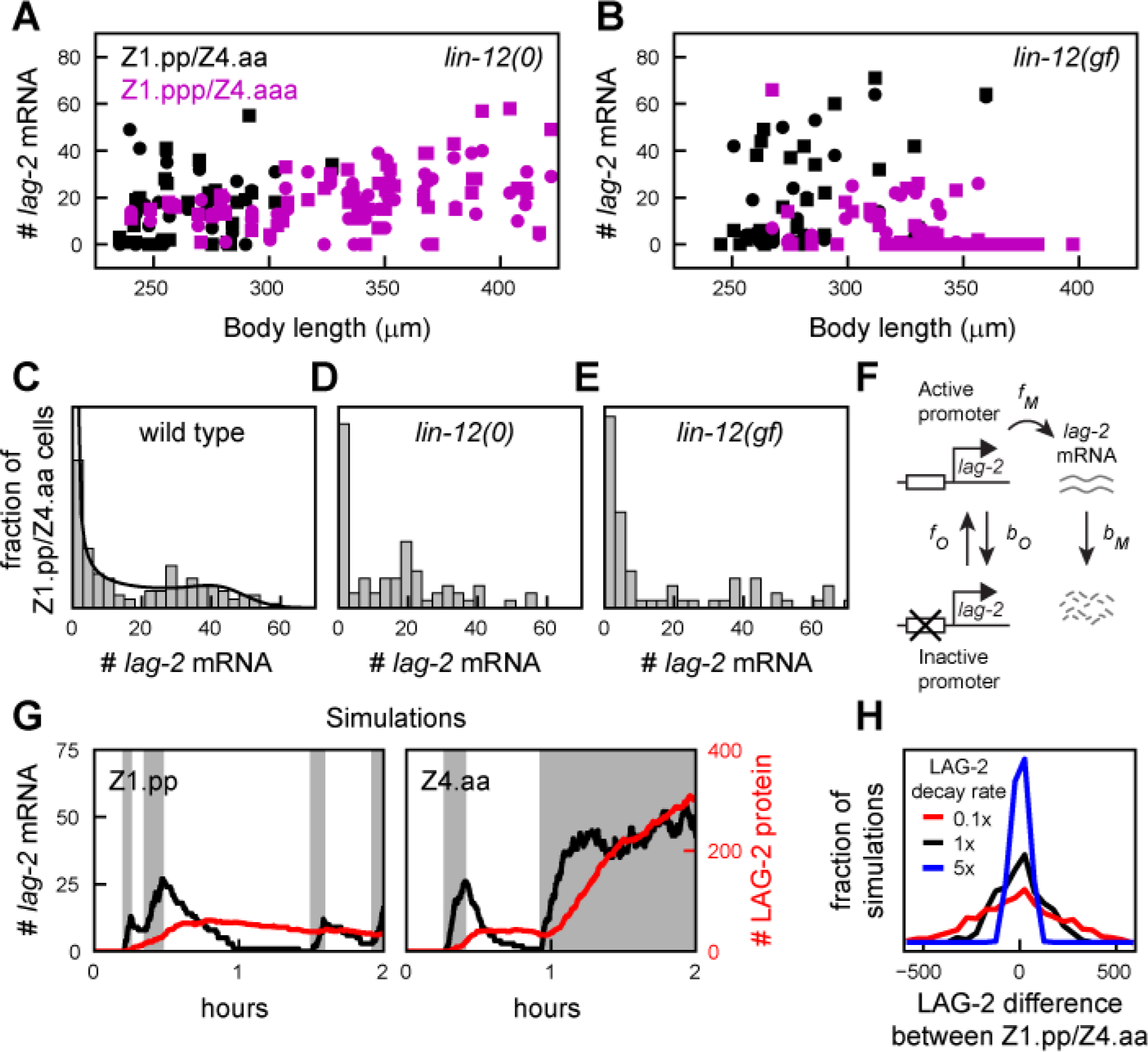
Bimodal variability in lag-2/Delta expression in Z1.pp and Z4.aa cells. **(a,b)** *lag-2* mRNA level versus body length in (a) *lin-12(0)* and (b) *lin-12(gf)* mutants. Black and magenta markers indicate Z1.pp/Z4.aa mother cells and Z1.ppp/Z4.aaa daughter cells, respectively, while circles and squares differentiate the Z1.pp/Z1.ppp and Z4.aa/Z4.aaa lineages. Lack of Notch signaling in *lin-12(0)* mutants causes both Z1.ppp/Z4.aaa daughter cells to express high *lag-2*, while hyperactive Notch signaling in *lin-12(gf)* mutants leads to inhibition of *lag-2* expression in those cells. **(c-e)** Distribution of *lag-2* mRNA levels in Z1.pp/Z4.aa mother cells in (c) wild-type, (d) *lin-12(0)* and (e) *lin-12(gf)* animals. All three genetic backgrounds show a bimodal *lag-2* mRNA distribution of *lag-2* expression, indicating that this shape of the distribution is independent of Notch signaling between Z1.pp and Z4.aa. The solid line in (c) is a fit to the model in (f). **(f)** Two-state model of *lag-2* expression. The *lag-2* promoter transitions stochastically between an inactive and active state, which *lag-2* mRNA transcription only in the latter. For sufficiently slow transitions, gene expression is ‘bursty’ and the model generates bimodal *lag-2* mRNA distributions (solid line in (c)). **(g)** Simulated *lag-2* mRNA (black) and protein (red) dynamics, resulting in strong stochastic differences in mRNA and protein level between Z1.pp and Z4.aa by the time of their division, ~2hr after birth. Grey intervals indicate time intervals when the *lag-2* promoter is in the active state. Simulations correspond to Eqs. 10-18 in the Methods. **(h)** Distribution of the difference in LAG-2 protein number between Z1.pp and Z4.aa, measured after 2hr, for LAG-2 protein degradation rate *b*_*D*_ = 0.1 hr^−1^(red), 1 hr^−1^(black) and 5 hr^−1^ (blue).

However, an alternative mechanism to explain such a bimodal distribution could be that one mother cell inhibits *lag-2* expression in the other by Notch signaling, analogous to the dynamics seen in their daughters Z1.ppp/Z4.aaa. Notch signaling is often assumed to start in Z1.ppp/Z4.aaa cells by the induction of *lag-2* expression upon their birth[20]. However, our observation that both the Notch ligand *lag-2* and the Notch receptor *lin-12* (Fig. 2c,d) are expressed already in the mother cells Z1.pp and Z4.aa, left open the possibility that Notch signaling already takes place between the Z1.pp and Z4.aa mother cell. However, we excluded that Notch signaling was responsible for the *lag-2* expression pattern observed in Z1.pp/Z4.aa mother cells by examining mutants that impacted Notch signaling. First, we found a similar bimodal *lag-2* distribution in a *lin-12* deletion mutant, *ok2215* (Fig. 4d), where in the absence of a functional Notch receptor *lag-2* remains expressed highly in the daughter cells Z1.ppp and Z4.aaa (Fig. 4a), causing both to assume AC fate[17]. Moreover, a similar *lag-2* distribution was also seen in the gain-of-function mutant *lin-12(gf)* (Fig. 4e). This is an even more striking result, because in this mutant the Notch receptor *lin-12* is activated even in the absence of Notch ligands, resulting in complete *lag-2* suppression in both Z1.ppp and Z4.aaa daughter cells (Fig. 4b), which subsequently assume VU fate[18]. The unperturbed *lag-2* expression in Z1.pp/Z4.aa in the *lin-12(gf)* mutant suggests that Notch signaling, or at least the inhibitory effect of Notch signaling on *lag-2* expression, is blocked in Z1.pp/Z4.aa cells and permitted only in their daughters after cell division.

In general, these results indicated that the bimodal distribution in *lag-2* expression was not due to Notch signaling, but rather cell-intrinsic stochastic *lag-2* expression. To gain insight in the underlying *lag-2* expression dynamics we fitted the experimentally observed *lag-2* mRNA distribution to a stochastic two-state model (Fig. 4f) used previously to study bursty gene expression[32,33]. Even though the model provided an excellent fit to the data (Fig. 4c, see Methods for fitting procedure), this result alone was not sufficient to provide information on the dynamics of *lag-2* expression, as it only provided the parameter ratios 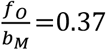, 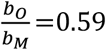 and 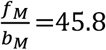, where *f*_0_ and *b*_0_ are the transition rates between the inactive and active *lag-2* promoter state and and are the *lag-2* mRNA production and degradation rate, respectively (Fig. 4f). However, we have previously measured the *lag-2* mRNA degradation rate directly in P6.p, a cell adjacent to the Z1.ppp/Z4.aaa cells, at roughly the same developmental stage as the AC/VU decision[34]. We used this measured rate in P6.p, *b*_*M*_ = 0.087 min^−1^, as an approximation. The resulting parameter values described a model where the *lag-2* promoter was in the actively describing state only *f*_0_/(*f*_0_+*b*_0_)=40% of the time. Together with the average expression burst duration 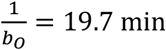, this corresponded to 2.4 bursts on average during the ~2 hour lifetime of the Z1.pp/Z4.aa cell. Additional model simulations highlighted that such a small number of bursts can give rise to strong stochastic differences in *lag-2* mRNA and protein number between the Z1.pp and Z4.aa mother cells (Fig. 4g), that become more pronounced for increased LAG-2 protein lifetime (Fig. 4h). Hence, bursty *lag-2* expression in the Z1.pp/Z4.aa mother cells could provide a strong noise source inherited by the Z1.ppp/Z4.aaa daughter cells to drive the AC/VU decision.

### Birth order bias in a mathematical model of the AC/VU decision

To understand the impact of birth order and stochastic *lag-2* expression on the AC/VU decision, we constructed a stochastic mathematical model of the underlying signaling network. The model (Eqs. 10-18 in Methods) has the following key ingredients: i) *lag-2* expression is bursty (Eqs. 11-14), ii) LAG-2 in one cell activates the Notch receptor LIN-12 in the other cell and iii) activated Notch signaling inhibits *lag-2* expression by inducing degradation of HLH-2, the transcriptional activator of *lag-2* (Eq. 10), as observed experimentally[20,21]. The model parameters governing *lag-2* expression were obtained by fitting the model to the *lag-2* mRNA distribution in the mother cells Z1.pp/Z4.aa (Fig. 4a) and the daughter cells Z1.ppp/Z4.aaa after the AC/VU decision was made (Supplementary Fig. 3). These fits show that *lag-2* expression is less bursty in the daughter cells Z1.ppp/Z4.aaa, compared to their mothers Z1.pp/Z4.aa. For LAG-2 protein, we assumed a protein life time of ~1hr and a copy number of ~500 molecules, with the resulting dynamics depending most strongly on the former. The remaining parameters were chosen so that the resulting model is bistable, with AC or VU fate assumed in a mutually exclusive fashion and driven by stochastic fluctuations (Fig. 5a, see Methods for full discussion of parameters).

**Figure 5.**
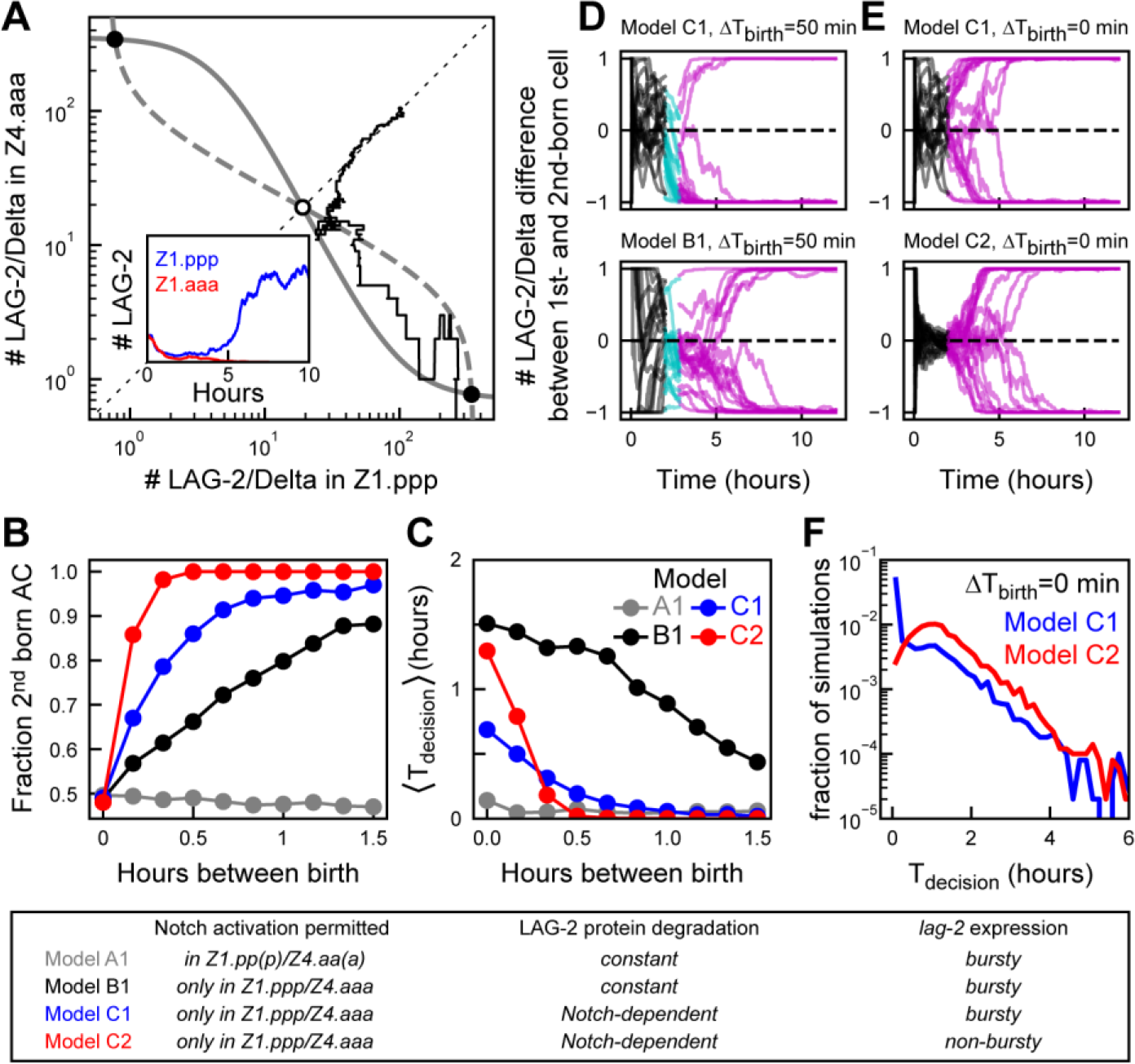
Birth order bias and decision times in a mathematical model of the AC/VU decision. **(a)** Symmetry breaking in a stochastic model of Notch signaling between Z1.ppp and Z4.aaa cells. Grey lines are nullclines of the differential equation model (Eqs. 1-9 in Methods) that predict two stable (black circle) and one unstable fixed point (white circle). The black line represents a stochastic simulation (Eqs. 10-18) with identical initial conditions for Z1.ppp and Z4.aaa and corresponds to the temporal dynamics shown in the inset. The simulation randomly results in *lag-2/Delta* expression restricted to Z1.ppp. **(b,c)** Comparing (b) birth order bias and (c) average time to decision 〈*T*_decision_〉 as a function of the time between Z1.ppp and Z4.aaa birth, Δ*T*_birth_, for the following models; A1, Notch activation in mother and daughter cells; B1, Notch activation only in daughter cells; C1, Notch-dependent LAG-2 degradation in daughter cells and; C2, as model C2 but with non-bursty *lag-2* expression in mother cells. For models A1 and B1, the fraction of second-born cells assuming AC fate as function of time between births is too low compared to the experiment in Fig. 3b. Model C1 is the best fit to the experimental data. Reducing variability in *lag-2* expression (model C2) causes exceptionally long decision times when Z1.ppp and Z4.aaa are born at similar times. For each Δ*T*_birth_, n=500 simulations. **(d,e)** Representative trajectories of simulations for the different models. Shown is the difference in LAG-2 protein number as function of time, with the difference defined as (*D*_1_-*D*_2_)/(*D*_1_ + *D*_2_) where *D*_1_ and *D*_2_ are the LAG-2 protein numbers in the first- and second-born daughter cell, respectively. Color indicates developmental stage; two mother cells (black), one cell (cyan) and both (magenta) cells divided. **(f)** Comparing the distribution of decision times for simultaneous Z1.ppp and Z4.aaa birth, for bursty (C1, blue) and non-bursty (C2, red) *lag-2* dynamics (*n*=5 ⋅ 10^3^ simulations). Model C1 shows more simulations with very short and fewer with very long decision times, compared to C2.

We used our experimental observations in Notch signaling mutants (Fig. 4) to constrain possible mechanisms underlying birth order bias. In the model, when Notch signaling was permitted both in the mother and daughter cells, Z1.pp/Z4aa and Z.1ppp/Z4.aaa, we observed no birth order bias, with equal probability of AC or VU fate independent of time between Z1.ppp/Z4.aaa birth (Fig. 5b, model A1). We found experimentally that activation of the Notch receptor LIN-12 did not cause inhibition of *lag-2* expression in mother cells Z1.pp/Z4.aa, but did so in daughter cells (Fig. 4b,e). Two mechanisms are consistent with these observations. First, that Notch signaling is only enabled between the daughter cells Z1.ppp/Z4.aaa. However, a model that incorporated this showed the opposite birth order bias, with the first-born cell preferentially assuming AC fate (Supplementary Fig. 4a, model A2). Second, Notch signaling cannot be activated in mother cells, but mother cells expressing *lag-2* can activate Notch signaling in an adjacent daughter cell. A model with these assumptions did show the correct birth order bias (Fig. 5b, model B1). Thus, our combined experimental and modeling results provide an attractive mechanism for birth order bias: a first-born daughter cell is biased towards VU fate, because it can receive an inhibitory Notch signal from the neighboring mother cell, while the mother cell is still insensitive to any inhibitory Notch signal from its neighbor.

### Notch-dependent LAG-2 degradation model reproduces birth-order bias and decision timescale

Even though model B1 exhibited birth-order bias, it was relatively weak: whereas we experimentally observed no first-born cells assuming AC fate after ~1 hour between births (Fig. 3b), ~20% of simulations had this outcome for model B1 (Fig. 5b,d). In such simulations, *lag-2* mRNA was rapidly down-regulated in the first-born cell, but LAG-2 protein levels remained high (Supplementary Fig. 4e) because its ~5-fold lower degradation rate compared to that of *lag-2* MRNA. This longer LAG-2 protein life-time in daughter cells means that more time is required for a mother cell to fully down-regulate LAG-2 in a neighboring daughter cell, thus leading to strong birth order bias only for long enough time between births.

One strategy for reducing LAG-2 protein life-time is increasing its turnover, i.e. increasing its production and degradation rate by the same factor both in mother and daughter cells. However, such a model showed weak birth order bias, even for long times between birth (Supplementary Fig. 4a, model B2). This occurred because increased protein turnover led to stronger fluctuations in LAG-2 number (Supplementary Fig. 4g) and, hence, in the level of inhibitory Notch signaling send to the neighboring cell. As a consequence, we often found LAG-2 levels rising, not falling, in the first-born cell, causing it to assume AC, rather than VU fate. Next, we increased the LAG-2 degradation rate specifically in daughter cells. We considered two models that achieved this: In this first model, C1, the LAG-2 degradation rate is increased in a Notch-dependent manner, i.e. with degradation rate increasing with strength of the incoming Notch signal (Eq. 19). In the second model, D1, the LAG-2 protein turn-over rate is increased in daughter cells compared to mother cells in a Notch-independent manner, by increasing both LAG-2 production and degradation rate in daughter cells by a factor 5. Both models reproduced the experimentally observed birth-order bias and increase of decision time for decreasing time between birth (Fig. 5b,c, Supplementary Fig. 4a,b). However, we found that the Notch-independent model D1 exhibited rare simulations with exceptionally long decision times, with *T*_dec_>6hr for 33/5000 simulations for D1, compared to 2/5000 for C1 (Supplementary Fig. 4c). Examining these simulations revealed that their dynamics differed significantly between models C1 and D1. For the model C1, all these simulations involved rare cases where both cells kept similar LAG-2 protein levels for many hours (Supplementary Fig 5h). However, for model D1, in most such simulations one cell initially emerged as the AC, but fluctuations in both cells destabilized this state, ultimately causing the other cell to assume AC fate (Supplementary Fig. 4h). This was due to increased variability in LAG-2 protein number, due to the increased LAG-2 protein turnover in both daughter cells. In model C1, however, LAG-2 degradation is Notch-dependent and, hence, high only in the presumptive VU cell, thereby minimizing fluctuations in the presumptive AC. For this reason, we choose C1 as the model that best fit the experimental data.

For all models examined, the decision time, *T*_dec_, was much shorter than experimentally observed (Figs. 3c, 5c). In general, for all models it was a challenge to generate long time to decision while simultaneously reproducing strong birth order bias for short, <30 min, time between birth. However, when we added to model C1 a long-lived YFP under control of a second, independent *lag-2* promoter (See Methods), comparable to the integrated *lag-2p::nls::yfp* transgene used in our experimental analysis, we found that the time to decision measured based on YFP level in the simulations had the same shape as that based on LAG-2 protein number, but with timescales closer to that observed experimentally (Supplementary Fig. 4d). This suggested that we likely overestimated the time to decision by measuring levels of long-lived YFP rather than short-lived LAG-2 protein. Overall, we found that a model, C1, where LAG-2 protein degradation rate in the daughter cells Z1.ppp/Z4.aaa is increased in proportion to the level of Notch signal received by cell, any symmetry between the two cells is broken in highly reliable manner, with birth order bias and the distribution of decision times as function of the time between births closely reproducing the experimental data.

### Bursty *lag-2* expression drives rapid decision when cells are born at similar times

Because it is difficult to experimentally change the variability in *lag-2* gene expression while keeping the average expression level unchanged, we instead used the mathematical model to assess role of bursty *lag-2* expression. We changed *lag-2* dynamics in the model with Notch-dependent LAG-2 degradation, C1, so that its average level was unchanged, but *lag-2* expression dynamics was no longer bursty (See Methods). We found that the resulting non-bursty model, C2, had significantly longer average time to decision when daughter cells are born close together in time, Δ*T*_birth_ ≈ 0 (Fig. 5c). Moreover, when comparing the full distribution for cells born simultaneously, we found that bursty *lag-2* gene expression strongly increased the fraction of animals with rapid decision time and decreased the fraction with very long decision time (Fig. 5f). This is because bursty *lag-2* expression decreases the fraction of animals where both daughter cells start the process of lateral Notch inhibition with similar LAG-2 protein levels, an initial condition that strongly correlates with long decision times (Supplementary Fig. 4i,j). However, despite the positive effect of bursty *lag-2* expression on reducing decision times for daughter cells born at similar times, it came at the expense of reducing birth order bias and increasing average decision times when daughter cells are born farther apart in time (Fig. 5b,c). This was because the resulting higher variability in LAG-2 protein levels increased the number of animals where LAG-2 levels in the first-born cell were significantly higher than average. This results in a substantial fraction of animals with similar LAG-2 levels in both daughter cells, leading to long decision times, or animals with higher LAG-2 level in the first-born cell, leading to rapid decisions but with the first-born cell assuming AC, not VU fate, even for large *T*_dec_ (Supplementary Fig. 4i,j). We found similar differences between for the model with increased LAG-2 protein turn-over in daughter cells, D1, and a variant of that model, D2, with non-bursty *lag-2* expression (Supplementary Fig. 4a-c), indicating that the effect of bursty *lag-2* expression is general.

If bursty *lag-2* expression is the main noise source if Z1.ppp/Z4.aaa are born at similar times, model C1 predicted that LAG-2 protein level in the mother cells Z1.pp/Z4.aa should correlate strongly with cell fate outcome, with the cell with highest LAG-2 levels biased strongly towards AC fate (Supplementary Fig. 5d). Even though in our experiments we measured strong variability in lag-2p::YFP levels between the mother cells Z1.pp/Z4.aa, we found no correlation between relative YFP levels in mother cells and cell fate outcome in animals where Z1.ppp/Z4.aaa were born within 30 mins of another (Supplementary Fig. 5a-c). However, we also observed no correlation between relative YFP level and outcome in model C1 (Supplementary Fig. 5e). In the model, this is because bursts in *lag-2* expression from the endogenous *lag-2* promoter are uncorrelated with those for *yfp* driven by the added *lag-2* promoter. Similarly, in the experiment any bursts in the integrated *lag-2* promoters driving *yfp* expression are likely uncorrelated with those of the endogenous *lag-2* promoter, explaining the observed lack of correlation. Hence, directly testing the link between bursty *lag-2* expression and cell fate outcome will require using a different transcriptional reporter than used here, namely one controlled by the native *lag-2* promoter. Such endogenous transcriptional reporter could be achieved experimentally, for instance using 2A peptides[35].

### *lag-2* levels impact the efficiency of the AC/VU decision

As the results above showed a critical role for changes in *lag-2* expression in restricting AC fate to a single cell, we tested the impact of changes in *lag-2* expression level or activity on the dynamics of the AC/VU decision. We first increased *lag-2* expression using a multi-copy integrated transgene (see Methods). Indeed, smFISH staining showed strongly elevated *lag-2* expression in the *lag-2(++)* overexpression mutant, with *lag-2* mRNA levels so high that individual molecules could no longer be discerned (Fig. 6a). Nevertheless, we found that *lag-2* expression became restricted to a single cell in all animals, both by smFISH (Fig. 6a) and by time-lapse imaging of *lag-2p::yfp::nls* dynamics (Fig. 6b), indicating that the AC/VU decision itself is robust to large changes in *lag-2* level. We then examined whether elevated *lag-2* expression also impacted the efficiency of the decision. First, we observed that the distribution of decision times was similar to that observed for wild-type animals (Fig. 6c), suggesting that any additional time required to break down the extra dose of *lag-2* mRNA and protein was still small compared to the slower timescale of YFP degradation. However, we found a significant difference between wild-type and *lag-2(++)* animals in birth order bias, with more first-born cells assuming AC rather than VU fate in *lag-2(++)* animals, even for cells born ~1 hour apart (Fig. 6d). In our model, we found that for efficient birth order bias *lag-2* mRNA and protein has to be fully degraded in the first-born cell before the other mother cell divides (Supplementary Fig. 4e,f). Hence, the weaker birth order bias in *lag-2(++)* animals likely reflected the longer time required to clear the elevated *lag-2* mRNA and protein levels from the first-born cell.

**Figure 6.**
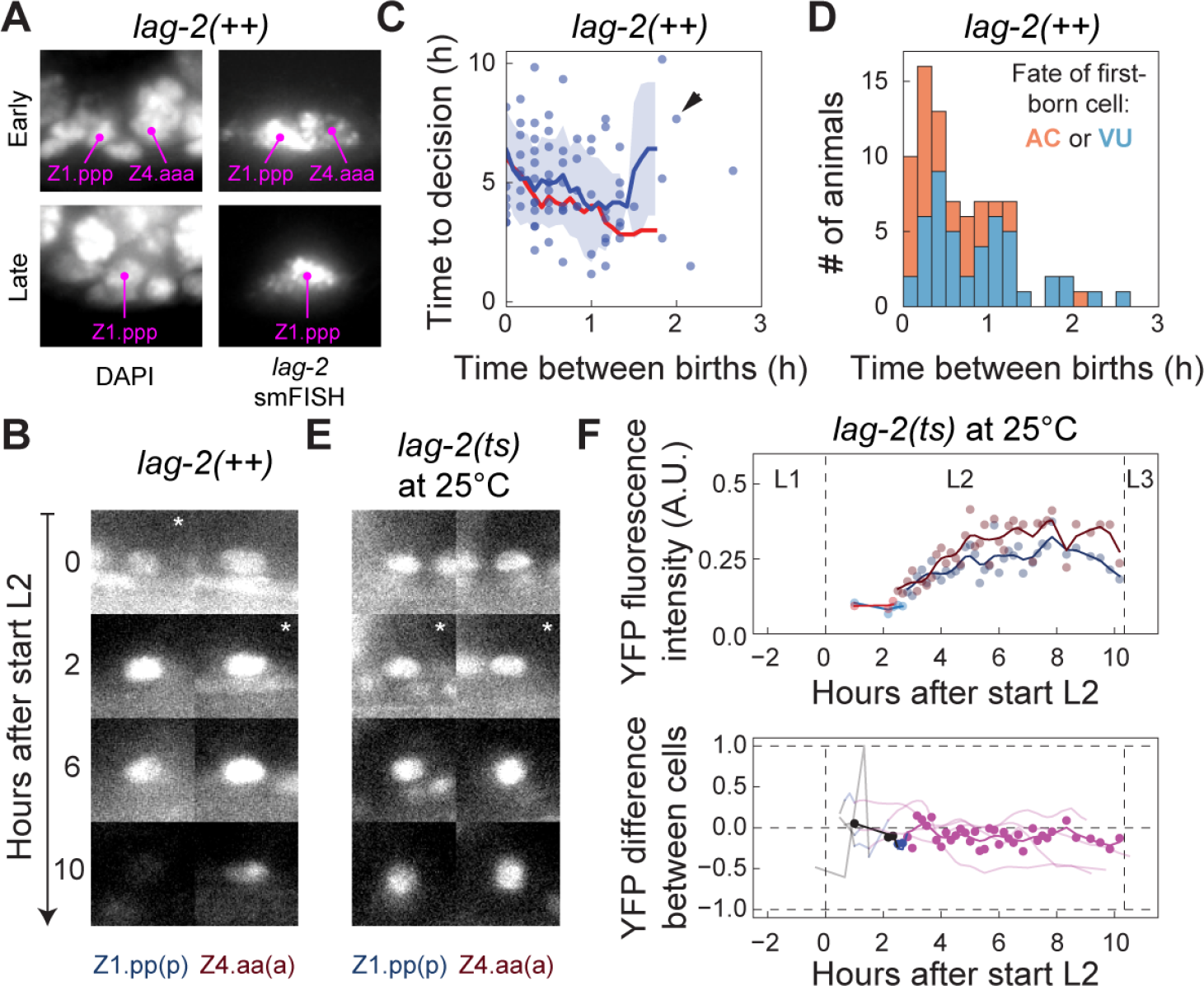
*lag-2*/Delta levels and activity impact AC/VU decision efficiency. **(a)** Increased *lag-2* expression in a *lag-2(++)* overexpression mutant at the early (top) and late (bottom) L2 stage. Levels are so high that individual mRNA molecules are no longer visible by smFISH. However, *lag-2* expression was still restricted to a single AC at late L2 (bottom). **(b)** Restriction of *lag-2p::nls::yfp* fluorescence to a single cell in a *lag-2(++)* animal. Starts indicate first time point after division. **(c)** Time to decision as function of time between birth of Z1.ppp and Z4.aaa, for the *lag-2(++)* mutant. Blue circles are individual animals. Shown is the average (blue line) and standard (shaded area). The average decision time and its dependence on time between births are similar to wild-type animals (red line). Marker corresponds to the animal in (b). **(d)** Birth order bias in the *lag-2(++)* mutant is weakened compared to wild-type animals (Fig. 3b), with the first-born cell assuming AC (orange) rather than VU fate (blue) in a significantly higher fraction of animals (two sample t-test with equal variance, P=0.0027). **(e)** In temperature-sensitive *lag-2* mutants grown at 25°C, reduced LAG-2 activity causes *lag-2p::nls::yfp* fluorescence to remain present in both Z1.ppp and Z4.aaa. **(f)** Even though YFP expression increases during L2 (top panel), no increase in difference between YFP levels in Z1.ppp and Z4.aaa is observed (bottom panel). Markers in both panels correspond to the animal in (e). Lines in bottom panel are dynamics in other animals.

Next, we studied the effect of decreasing *lag-2* activity on AC/VU decision dynamics. For this, we used a temperature-sensitive *lag-2* loss-of-function mutant, *lag-2(q420)*, that was previously shown to exhibit 2 ACs when grown at 25°C[17,20,36]. Because loss of *lag-2* is lethal during early development[17], we started our time-lapse microscopy at 20°C and shifted to 25°C after all animals has reached the L2 larval stage. However, under this treatment many animals still arrested development, severely limiting the number of animals we could examine. Nevertheless, in those animals where we could observe the AC/VU decision, we saw a striking lack of symmetry-breaking in *lag-2p::nls::yfp* expression between the Z1.pp(p) and Z4.aa(a) cells (Fig. 6e,f). In general, most animals maintained similar YFP levels throughout the L2 larval stage (Fig. 6g). Some animals showed a reduction in YFP fluorescence in one daughter cell, but with a much smaller difference between Z1.ppp and Z4.aaa at the end of the L2 stage than observed in wild-type.

## Discussion

Here, we studied the dynamics of the AC/VU decision, a stochastic cell fate decision based on lateral Notch inhibition. Notch-mediated stochastic cell fate decisions are omnipresent in development[37], but it has remained an open question how small stochastic differences between cells, i.e. ‘noise’, is amplified by Notch signaling into differences in cell fate. To address this question, we quantified the expression dynamics of the Notch ligand *lag-2/Delta* and the Notch receptor *lin-12/Notch* using time-lapse microscopy (Fig. 1) and single molecule FISH (Fig. 2), and used the observed dynamics to constrain mathematical models of the AC/VU decision (Fig. 3-5).

Surprisingly, we found that both Notch ligands and receptors were already expressed in Z1.pp and Z4.aa cells, even though the restriction of AC and VU fate to a single cell only occurred in their daughter cells Z1.ppp and Z4.aaa (Fig. 2). Another surprising observation was that high *lin-12* mRNA (Fig. 2) and protein levels (Supplementary Fig. 2) were not restricted to a single cell even at the end of the L3 larval stage, when *lag-2* expression was already restricted to either Z1.ppp and Z4.aaa. So far, it is often considered that symmetry breaking by Notch signaling requires Delta ligands and Notch receptors to change their gene expression in a reciprocal manner, with Notch activation leading both to higher Delta ligand and lower Notch receptor expression. Our experimental observations show that robust Notch-mediated symmetry breaking can be achieved by changing expression only of Delta ligands, with the Notch receptor serving merely as passive communication channel. Indeed, this notion is supported by our mathematical model, which showed bistability (Fig. 5a) even when assuming constant *lin-12/Notch* levels (Eq. 7). In the model, this requires that degradation of HLH-2, the activator of *lag-2/Delta* expression, depends on Notch activation in a sufficiently cooperative manner (Eq. 7).

It was observed previously that the Z1.ppp and Z4.aaa are born with variable timing and the resulting random birth order provides a strong bias to the AC/VU decision, with the first-born cell biased towards VU fate[20]. Because of our ability to follow the AC/VU decision in live animals by time-lapse microscopy we could study the impact of birth order on cell fate outcome with substantially higher throughput, and also investigate for the first time how birth order and the time between the birth of Z1.ppp/Z4.aaa impacted the expression dynamics of the *lag-2* promoter. We found that birth order correlated strongly with cell fate when Z1.ppp/Z4.aaa were born more than 1 hour apart (Fig. 3b). For animals with shorter time between births, we not only found more frequently that first-born cells assumed AC, not VU fate, but also observed that more time was required for the expression of the transcriptional *lag-2* promoter to be restricted to a single cell (Fig. 3c). Overall, this identified birth order bias as a key noise source driving the AC/VU decision in animals where Z1.ppp/Z4.aaa were born sufficiently far apart in time, but raised the question what noise source drove the decision in animals where these cells were born at similar times.

It remains an open question how birth order impacts Notch signaling to bias cell fate outcome[20]. Here, we showed that the mother cells Z1.pp/Z4.aa and daughter cells Z1.ppp/Z4.aaa respond to loss or ectopic activation of Notch signaling in a strikingly different manner: whereas a *lin-12/Notch* null mutant and a constitutively active *lin-12/Notch* mutant in either high *lag-2/Delta* expression or complete absence of *lag-2/Delta* expression in the Z1.ppp/Z4.aaa daughter cells, we found that *lag-2/Delta* expression in their mother cells Z1.pp/Z4.aa was not impacted in either mutant (Fig. 4). This suggested that Z1.pp/Z4.aa cells are not able respond to Notch activation by changing *lag-2/Delta* expression, but do so immediately after they divide. This is consistent with previous results that show that Notch activation during *C. elegans* vulva induction is strongly cell cycle dependent, with Notch signaling inhibited in the pre-mitotic G2 phase of the cell cycle [38,39]. Indeed, given the short, ~2hr, cell cycle duration we observed in Z1.pp/Z4.aa, it is conceivable that they exist mostly in S and G2, rather than G1 phase. This asymmetry in Notch signaling between mother and daughter cells means that the first-born cell is impacted by any Notch signal from the adjacent mother cell while the reverse is not the case, providing a clear molecular mechanism for birth order bias. Indeed, only when we assumed in our model that mother cells are able to send but not receive a Notch signal, but daughter cells are capable of both, did we observe the correct dependence of first-born cell fate on birth order (Fig. 5b).

In the experiments, birth order biased cell fate outcome already when cells were born ~30 minutes apart in time, even though fluorescence from the *lag-2/Delta* transcriptional reporter required a significantly longer time to become restricted to a single cell (Fig. 3b,c). In the model, we could only reproduce the short timescale over which birth order bias becomes dominant by assuming that LAG-2/Delta protein degradation was rapid (Fig. 5b,c), with a ~10 min half-life comparable to *lag-2/Delta* mRNA and much faster than the rate of YFP degradation observed in the *lag-2/Delta* reporter (Supplementary Fig. 4d). In the model for slow LAG-2/Delta protein degradation, there was insufficient time for all Notch ligands to be cleared from the first-born cell if the other cell was born within ~30 minutes, causing a significant fraction of first-born cells to assume AC fate (Supplementary Fig. 4e,f). In our model, we found that the stability of the AC/VU decision was optimal when the rate of LAG-2 protein degradation increased with Notch activity (Fig. 5f, Supplementary Fig. 4h), because in this case LAG-2/Delta was rapidly degraded in the prospective VU cell but remained stable in the AC. It is known that Notch activation causes degradation of HLH-2 [20,21], the transcriptional activator of *lag-2/Delta* expression, but there is currently no evidence linking it to LAG-2/Delta degradation. In general, the optimal LAG-2/Delta protein degradation rate in the model was fast compared to the typical protein life time of many hours[40]. However, whereas in the model we assumed high LAG-2 protein degradation, the same result could be obtained by other mechanisms, e.g. LAG-2 endocytosis, that reduce the amount of ligand available for Notch signalling. The importance of rapid LAG-2 degradation for strong birth order bias was underscored by our observation that in animals overexpressing *lag-2/Delta* birth order bias was significantly weakened, with first-born cells assuming AC fate in animals where Z1.ppp/Z4.aaa were born 1-2 hours apart (Fig. 6).

Surprisingly, we found that *lag-2* expression in the mother cells Z1.pp/Z4.aa was highly variable, showing a bimodal distribution that was independent of Notch signaling (Fig. 4). This suggested that transcription of *lag-2/Delta* expression in Z1.pp/Z4.aa occurred in bursts. Indeed, the observed bimodal distribution could be fitted well with a bursty gene expression model[33] that assumed that the *lag-2/Delta* promoter exhibited slow, stochastic transitions between a transcriptionally active and inactive state (Fig. 4c,f). The fitted values of the model parameters, together with an estimated ~10 min half-life for *lag-2/Delta* mRNA, indicated that only 1-5 transcriptional bursts occurred in the ~2 hour lifetime of Z1.pp/Z4.aa cells and these could lead to substantial variability in *lag-2/Delta* mRNA and protein level between Z1.pp/Z4.aa prior to their division (Fig. 4g,h). Indeed, by comparing models with and without bursty *lag-2/Delta* expression, we found that this increased variability between cells in the model with bursty *lag-2/Delta* expression resulted in much reduced decision times specifically for simulations where Z1.ppp and Z4.aaa were born close together in time (Fig. 5,c,f). These results showed that bursty *lag-2/Delta* expression is likely a key alternative noise source driving the AC/VU decision in animals where cells are born close together in time and birth order bias forms a weak noise source.

However, this beneficial effect of bursty *lag-2/Delta* expression appeared to come at a cost: whereas it stimulated short decision times when Z1.ppp/Z4.aaa were born close together in time, compared to a model with non-bursty *lag-2/Delta* expression it showed an overall weaker birth order bias and longer decision times when cells were born far apart in time (Fig. 5b,c). In the model, this occurred when the two noise sources interfered with each other’s effect, e.g. when bursty *lag-2/Delta* expression resulted in high *lag-2/Delta* levels in the first-born but low levels in the second-born cell, requiring a longer time for the second-born cell to repress *lag-2/Delta* in its neighbor. In *C. elegans*, both the AC and VU cells are required for subsequent developed, with the AC required for vulva induction and morphogenesis from the late-L2 stage onwards and the VU cells dividing again in the early-L3 stage. This means that both cells have to be specified in a 5-10 hour time window during the L2 stage. Our observation of bursty *lag-2/Delta* expression in the Z1.pp/Z4.aa mother cells in the late-L1/early-L2 stage indicated that for development it is more important to prevent cases where the AC/VU decision last exceptionally long in animals where Z1.ppp/Z4.aaa are born close together in time than to reduce the average time of the decision for all animals. Unpredictability is a fundamental property of stochastic cell fate decisions that seems at odds with the clockwork precision of development. It will be interesting to examine whether mechanisms that prevent extreme fluctuations in the dynamics of stochastic cell fate decisions, even at the expense of average efficiency, occur more generally in (Notch-based) stochastic cell fate decisions.

## Materials and Methods

### *C. elegans* strains and culture

All strains were handled according to standard protocol [41]. Wild-type nematodes were strain N2.

### Genetics

The following transgenes were used: *arIS131[lag-2p::2xNLS::YFP]*[29], *itIS37[pie-1p::mCherry::H2B::pie-1]*, *wgIs72 [lin-12::TY1::EGFP::3xFLAG(92C12) + unc-119(+)]* [31]. The following mutant was used to perform the temperature sensitive experiment: *LGV: lag-2(q420)*. To perform the *lin-12(0)* experiments, the mutant *lin-12(ok2215) III/hT2 [bli-4(e937) let-?(q782) qIs48]*[42] was used. For the *lin-12(gf)* experiments the mutant *LGIII: lin-12(n302)* was used.

### Transgenesis

To increase the number of lag-2 copies, multi copies insertion animals were generated. The *lag-2* multi-copy integrated line was created by injecting a *lag-2* full-length construct, consisting of a 7106 bp long promoter region directly upstream of the translational ATG site of *lag-2*, the entire 1492bp *lag-2* coding region, and a 1460bp long 3’ UTR of *lag-2*, into the MosScI insertion line EG6699 (*ttTi5605)*. To create the *lag-2* full-length construct we used the Multisite-Gateway Three-Fragment Vector construction kit (Thermo Fisher Scientific, 2012). 60ng /µl of this 17.623bp long *lag-2* full-length construct was injected together with the co-injection markers (pCFJ601 50ng/µl, pmA122 10ng/µl, pGH8 10ng/µl, pCFJ90 2,5ng/µl and pCFJ104 5ng/µl) into EG6699 animals using standard injection techniques[43]. To integrate the extrachromosomal *lag-2* transgene we used gamma irradiation, following the standard technique[43].

### Time-lapse microscopy

Time-lapse imaging was performed as previously described[28]. Briefly, microfabricated arrays of chambers were made in polyacrylamide hydrogel using standard soft lithography techniques. We filled each of these chambers with OP50 bacteria and a single *C. elegans* embryo. We used chambers of 190×190×10μm for all time-lapse experiments, as we found that chambers of this dimension contain sufficient OP50 bacteria to sustain development until well after the AC/VU decision has occurred.

Time-lapse imaging was performed on a Nikon Ti-E inverted microscope using a 60X magnification objective (Nikon Plan Fluor 60X, N.A. 1.4, oil immersion). We used the Hamamatsu Orca Flash 4.0 v2 camera at full frame and full speed. The camera chip is 13×13mm in size, therefore we could achieve a field of view of ~220μm, sufficient to accommodate the microfabricated chambers in the field of view of a single image. At the same time, the camera chip contains 4Mp, therefore each pixel collects light from a 100×100nm region in the sample, which is sufficient to resolve single cells in a developing *C. elegans* larva. We used 561nm and 488 nm lasers (Coherent OBIS-LS 488-100 and Coherent OBIS-LS 561-100) for fluorescence excitation. Because of the bright fluorescence, we found that high signal to noise ratio was always achieved with a laser power of approximately 10 mW and an exposure time of 10 ms. Bright field imaging was performed with a red LED (CoolLED, 630nm) to minimize photophobicity effects. At maximum speed, a full Z-stack containing 20 focal planes in 3 different channels (bright field, 488nm, 561nm) was acquired in ~1 second. Time-lapse images were acquired every 10 minutes without detectable photoxicity effects.

### Image analysis

To analyze the large amount of data created, we developed custom written software (available at https://github.com/Nikoula86/ACVU_timelapse_analysis). Every step in the image analysis was performed using a script optimized for the specific task. First, a maximum intensity projection movie of every acquired channel was generated and the timepoints at which the worm hatched, entered L2 and L3 stages were annotated. Because of the short time necessary to acquire a single stack, the animals were found in the same position throughout all the frames in a stack. Therefore, the (x,y) position of the gonad at every timepoint was manually annotated based on the maximum intensity projection image only and this information was used to crop a 512×512pixel=55×55μm area around the center of the gonad from the full 3D stack. While all the ~60 Gb of raw data were saved and stored in a separate disk, this step allowed us to reduce the data containing useful information for the subsequent analysis to ~4 Gb of local disk space per animal. Next, the cropped stacks were visually inspected and the cell position were manually detected and labelled. All the images for a single animal could be annotated in ~15 minutes. Importantly, at the end of each experiment, we acquired a flat field, F, and a dark field, D, image and corrected the raw image R on a pixel by pixel basis, according to the 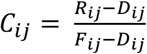. All subsequent analysis was performed on the corrected image C.

To generate all images and movies, we annotated the anterior, posterior and dorsal side of the animal around the gonad, and used this reference points to orient the images such that the head always lays on the left, tail to the right and dorsal side to the top of the image. For quantification purposes, we computed the average fluorescence in a circular area of ~5μm diameter around every annotated cell position and corrected this value by subtracting the background level found around each cell. We then determined the fluorescence intensity ratio as 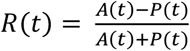 where *A*(*t*) and *P*(*t*) represent the fluorescence intensity of the anterior and posterior cells, respectively. We then annotated each cells as Z1.pp, Z1.ppp, Z4.aa or Z4.aaa, depending on the observed division time of each mother cell. Fluorescence trajectories were obtained by convolution of the fluorescence time-series *R*(*t*) with a Gaussian function with standard deviation of 20 minutes.

### Single molecule FISH

To visualize mRNA transcripts in L1 and L2 larvae (5-16 hours after hatching), probe design and smFISH hybridization were performed as described previously[24,25]. Briefly, animals were collected by washing plates with dH2O for two times (to remove bacteria and agar) and were fixed in 4% formaldehyde in 1XPBS for 45 min at room temperature. Fixed animals were permeabilized in 70% ethanol overnight at 4°C. Probes for smFISH hybridization were coupled to Cy5 (GE Amersham) or Alexa594 (Invitrogen). Images were acquired with a Nikon Ti-E inverted fluorescence microscope, equipped with a 100X plan-apochromat oil-immersion objective and a Princeton Instruments Pixis 1024 CCD camera controlled by MetaMorph software (Molecular Devices, Downington, PA, USA). In addition, nuclei were visualized with DAPI for cell identification. Exact three-dimensional positions of smFISH fluorescent spots in each animal were detected using a custom MATLAB (The Mathworks) script, based on a previously published algorithm[25].

### Differential equation model of the AC/VU decision

#### Promoter dynamics

In our model, the *lag-2* promoter can exist in four configurations (Supplementary Fig. 6): an open (*O*) and closed (*O*\*) configuration without HLH-2 bound and an open (*OH*) and closed (*O*H*) configuration with HLH-2 bound. The rate of transitions between the open and closed configuration, *f*_O_ and *b*_O_, are independent of whether HLH-2 is bound. Similarly, the rates of HLH-2 (un)binding to the *lag-2* promoter, *f*_OH_ and *b*_OH_, are independent of whether it is in the open or closed configuration. This leads to the following differential equations for the levels of the four promoter configurations in cell *i*:

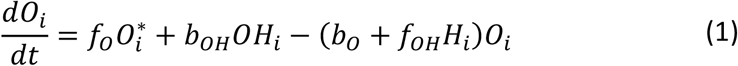

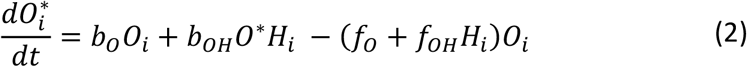

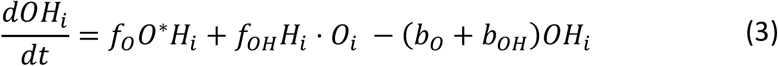

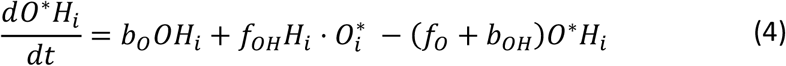

#### LAG-2 mRNA and protein production

We assume that *lag-2* is transcribed only when HLH-2 is bound to the open configuration (Supplementary Fig. 6) leading to the following differential equations for the *lag-2* mRNA level *M*_*i*_ and LAG-2 protein level *D*_*i*_:

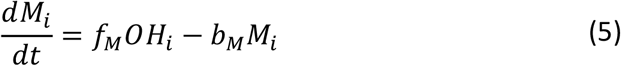

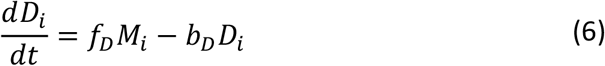

#### HLH-2 dynamics

We assume that HLH-2 is continuously produced at rate *f*_*H*_ and degraded at rate *b*_*H*_ independent of Notch signaling. However, in response to a Notch signal *S*_*ij*_ received by cell *i* from neighboring cell *j*, HLH-2 in cell *i* is degraded at a higher rate that depends on the strength of the Notch signal in non-linear manner:

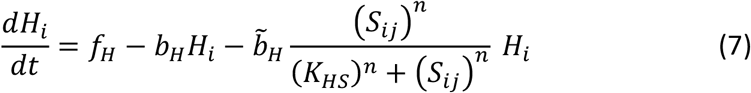

We assumed that the strength of the Notch signal was given by *S*_*ij*_ = ϕ_*i*_*D*_*i*_, where *D*_*j*_ is the Notch ligand level in neighboring cell *j* and ϕ_*j*_ is a coupling constant that describes how strongly a given Notch signal is received by cell *i*.

#### Steady state two-cell solution

To examine for what parameter values our model exhibited bistable behavior, we calculated the steady state solution of Eqs. 1-7. We found an analytical expression of the configuration with HLH-2 bound to the open complex, a state associated with high *lag-2*/Delta expression:

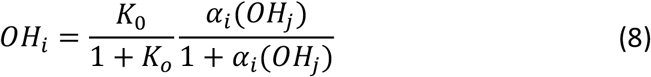

where

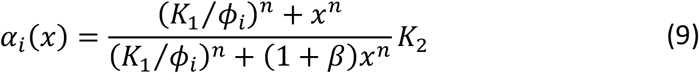

Here,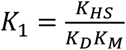 and*K*_2_ = *K*_*OH*_*K*_*H*_, with the equilibrium constants given by *K*_*O*_ = *f*_*O*_/*b*_*O*_, *K*_*H*_ = *f*_*H*_/*b*_*H*_, *K*_*OH*_ = *f*_*OH*_/*b*_*OH*_, *K*_*M*_ = *f*_*M*_/*b*_*M*_ and *K*_*D*_ = *f*_*D*_/*b*_*D*_. Finally, the parameter 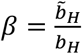 is the ratio between the rate of Notch-dependent and spontaneous HLH-2 degradation. For a system with two cells, fixed points occur where the nullclines *OH*_1_ and *OH*_2_ intersect (Fig. 5a). For sufficiently cooperative degradation of HLH-2 by Notch activation, *n*=2, the model exhibits one unstable and two stable fixed points, a hallmark of bistability (Fig. 5a).

### Stochastic model of the AC/VU decision

#### Basic model

We assume that HLH-2 production and degradation dynamics is fast compared to all other reactions. In that case, the rate of HLH-2 binding to the *lag-2* promoter is given by the effective rate 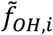:

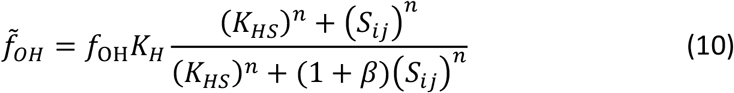

where *S*_*ij*_ = ϕ_*i*_*D*_*j*_. With this assumption, the set of reactions describing the differential equation model in Eqs. 1-7 is given by:

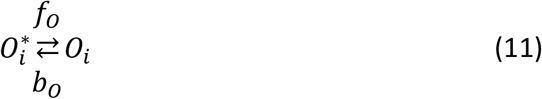

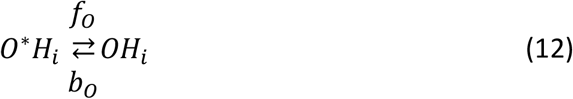

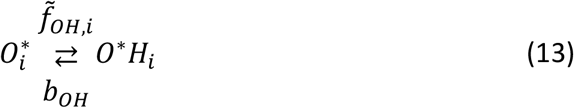

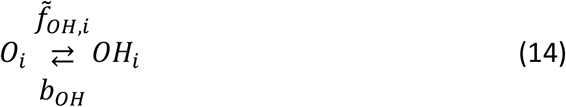

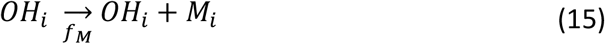

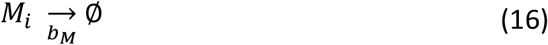

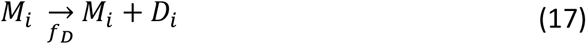

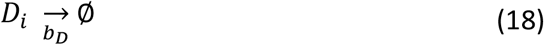

For Model A1, where both mother cells Z1.pp/Z4.aa and daughter cells Z1.ppp/Z4.aaa interact by Notch signaling, we set the Notch coupling constant to ϕ_*i*_ = 1 for both mother and daughter cells. We also examined a model (Model A2) where Notch signaling only occurs between daughter cells. In this case, ϕ_*i*_ = 1 only when both daughter cells Z1.ppp and Z4.aaa are present and ϕ_*i*_ = 0 for all other combinations of mother and daughter cells. For all other models, daughter cells do, but mother cells do not respond to an incoming Notch signal from the adjacent cell. In this case, ϕ_*i*_ = 1 if cell *i* is a daughter cell, i.e. Z1.ppp/Z4.aaa, and to ϕ_*i*_ = 0 if cell *i* is a mother cell, i.e. Z1.pp/Z4.aa.

For models B1 and B2, the LAG-2 protein degradation rate has the same low (Model B1) or high (Model B2) value both in mother and daughter cells. For Models D1 and D2, the rate of LAG-2 protein turn-over was increased in the daughter cells Z1.ppp/Z4.aaa compared to their mothers Z1.pp/Z4.aa. Model D2 is the same as Model D1, except that *lag-2* expression is not bursty. To do so, we removed the closed promoter configuration *O*\*, by setting the rate of transition to the closed configuration to *b*_*O*_ = 0. Moreover, we changed the mRNA production rate *f*_*M*_ so that the average mRNA level is the same as in Model D1. Specifically, we set 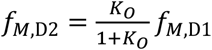, where *K*_*O*_ = *f*_*O,D1*_/*b*_*O,D1*_.

#### Model with Notch-induced LAG-2/Delta protein degradation

For Models A, B and D, downregulation of LAG-2 only occurs by inhibition of *lag-2* expression, with LAG-2 decay occuring at the spontaneous LAG-2 degradation rate *b*_*D*_ (Eq. 18). For Models C1 and C2, we assumed that Notch receptor activation leads to degradation of LAG-2 protein with a higher degradation rate. We implemented this by replacing the constant LAG-2 degradation rate *b*_*D*_ in Eq. 18, with a rate that depends on the degree of Notch receptor activation:

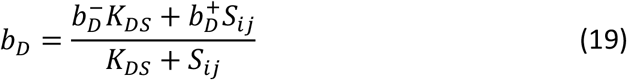

where 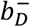 is the degradation rate in the absence of Notch signaling and the degradation rate for strong Notch signaling, i.e. *S*_*ij*_ ≫ *K*_*DS*_, is given by 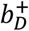. Both models differ in whether *lag-2* mRNA production is bursty (Model C1) or not (Model C2). Parameters for Model C2 were adjusted from those for Model C1, in the same way as described above for Model D2.

#### Incorporating lag-2p::yfp::nls dynamics

To include YFP expression from a second, independent lag-2 promoter, similar to the lag-2p::yfp::nls reporter strain, we assumed that in this case yfp mRNA is produced from an operator that can transition between a empty state, *O*_*Y*_, and one bound by HLH-2, *O*_*Y*_*H*, with forward rate *f*_*O_Y_H*_ ⋅ *H* and backward rate *b*_*O_Y_H*_, where *H* is the steady state HLH-2 level calculated from Eq. 7. This is similar to the dynamics in Eqs. 11-14, but here for simplicity we assume *yfp* expression is not bursty and hence we do not consider a closed state. YFP expression is modeled similar to Eqs. 15-18: mRNA transcription only occurs from the *O*_*Y*_*H* promoter state, meaning that HLH-2 is bound to the promoter, with rate *f*_*MY*_ and mRNA degradation rate *b*_*MY*_ and YFP protein translation occurs with production rate *f*_*Y*_ and degradation rate *b*_*Y*_.

### Simulation and parameters

To study the dynamics of the different models, we performed stochastic Gillespie simulations [44]. We used a custom written Python script to simulate reactions in the two cells, for all possible combinations of cells: two mother cells, one daughter and one mother cell and two daughter cells. The full simulation is designed as follows: 1) execute the reactions in Eqs. 11-18 with mother cell-specific parameters for a time *T*_0_ =2 hr, 2) divide one mother cell to generate the first-born daughter cell, 3) execute the reactions with parameters specific for a mother cell-daughter cell pair for time Δ*T*_birth_, 5) divide the remaining mother cell, and 6) execute the reactions with daughter cell-specific parameters for a time *T*_1_ − Δ*T*_birth_Cell divisions were implemented as follows: upon division, *lag-2* mRNA (*M*) and protein (*D*) present in the mother cell, e.g. Z1.pp, were assumed to be randomly inherited by the daughter cells, e.g. Z1.ppa and Z1.ppp, with equal probability. This was implemented by randomly drawing a new copy number *n*_*i*_ for each chemical species *i* = *M, D* from a binomial distribution 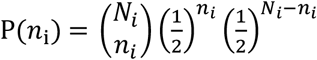, where *N*_*i*_ is the copy number in the mother cell upon the time of division. For the remaining chemical species describing the *lag-2* promoter state, *O*, *O*\*, *OH*, *O*H*, we assumed that daughter cells inherited the promoter state from their mother. Simulations are initiated without *lag-2* mRNA and protein and the *lag-2* promoter unbound by HLH-2 and either in the closed (Models A1-2, B1-2, C1 and D1) or the open (Models C2 and D2) configuration.

We choose parameter values as follows: we based the *lag-2* mRNA degradation rate on direct measurements we performed in another cell (P6.p) at a similar stage of development[34]. The mother and daughter cell specific-parameters *f*_*O*_, *b*_*O*_, *f*_*M*_ were obtained by fitting the experimentally observed *lag-2* mRNA distribution (Fig. 4c and Supplementary Fig. 3). The rates *f*_*OH*_, *b*_*OH*_ and *k*_*H*_ were chosen so that HLH-2 binding to the *lag-2* promoter was fast compared to the other reactions and, when present, bound strongly to the *lag-2* promoter. The parameters *K*_*HS*_, *n* and *β* were chosen so that the model was bistable (Fig. 5a) with almost complete inhibition of HLH-2 levels in response to Notch signaling. There exist no fluorescently labeled translational LAG-2 fusions and, as a consequence, there is no good estimate for LAG-2 protein copy number. The LAG-2 protein production and degradation rates *f*_*D*_ and 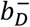 were chosen that that the average LAG-2 protein copy number in the AC was ~350 molecules. For production of *yfp* mRNA and protein, we used the same values as for *lag-2*. In the table below, we summarize all parameter values. In this table, all parameters values are defined for the best fit model, C1, and for the other models parameters values are only given when they deviate from those used for Model 1.

| Parameter | Description | Value |
| --- | --- | --- |
| **Model C1** | **Notch-dependent LAG-2 protein degradation rate, Notch signaling permitted between mother and daughter cells (best fit model)** |  |
| $f_O$ | Transition rate to open promoter configuration **in Z1.pp/Z4.aa** | $0.033 \text{ min}^{-1}$ |
| | Transition rate to open promoter configuration **in Z1.ppp/Z4.aaa** | $0.42 \text{ min}^{-1}$ |
| $b_O$ | Transition rate to closed promoter configuration **in Z1.pp/Z4.aa** | $0.051 \text{ min}^{-1}$ |
| | Transition rate to closed promoter configuration **in Z1.ppp/Z4.aaa** | $0.43 \text{ min}^{-1}$ |
| $f_M$ | *lag-2* mRNA production rate **in Z1.pp/Z4.aa** | $3.99 \text{ min}^{-1}$ |
| | *lag-2* mRNA production rate **in Z1.ppp/Z4.aaa** | $6.32 \text{ min}^{-1}$ |
| $b_M$ | *lag-2* mRNA degradation rate | $0.087 \text{ min}^{-1}$ |
| $f_D$ | LAG-2 protein production rate | $0.17 \text{ min}^{-1}$ |
| $b_D^-$ | Spontaneous LAG-2 protein degradation | $0.017 \text{ min}^{-1}$ |
| $b_D^+$ | Notch-induced LAG-2 protein degradation | $0.08 \text{ min}^{-1}$ |
| $K_{DS}$ | Threshold for Notch-induced LAG-2 protein degradation | $1 \cdot 10^2$ |
| $f_{OH}$ | Binding of HLH-2 to *lag-2* promoter | $1 \cdot 10^3 \text{ min}^{-1}$ |
| $b_{OH}$ | Unbinding of HLH-2 from *lag-2* promoter | $1 \cdot 10^2 \text{ min}^{-1}$ |
| $K_H$ | Equilibrium constant for HLH-2 production/degradation in absence of Notch signaling | 10 |
| $K_{HS}$ | Threshold for degradation of HLH-2 in response to Notch activation | 100 |
| $n$ | Hill coefficient for Notch induced HLH-2 degradation | 2 |
| $\beta$ | ratio between Notch-dependent and spontaneous HLH-2 degradation rate | $5 \cdot 10^4$ |
| $\phi$ | Notch coupling constant **in Z1.pp and Z4.aa** | 0 |
|  | Notch coupling constant **in Z1.ppp and Z4.aaa** | 1 |
| **Model A1** | **Constant LAG-2 protein degradation, Notch signaling permitted between mother and daughter cells** |  |
| $b_D$ | LAG-2 protein degradation rate | $0.017 \text{ min}^{-1}$ |
| $\phi$ | Notch coupling constant for all cells | 1 |
| **Model A2** | **Constant LAG-2 protein degradation, Notch signaling permitted only between daughter cells** |  |
| $b_D$ | LAG-2 protein degradation rate | $0.017 \text{ min}^{-1}$ |
| $\phi$ | Notch coupling constant **between Z1.ppp and Z4.aaa** | 1 |
|  | Notch coupling constant **for all other cell combinations** | 0 |
| **Model B1** | **Constant LAG-2 protein degradation** |  |
| $b_D$ | LAG-2 protein degradation rate | $0.017 \text{ min}^{-1}$ |
| **Model B2** | **Constant high LAG-2 protein degradation** |  |
| $b_D$ | LAG-2 protein degradation rate | $0.08 \text{ min}^{-1}$ |
| **Model C2** | **Non-bursty *lag-2* expression** |  |
| $b_O$ | Transition rate to closed promoter configuration **in Z1.pp(p)/Z4.aa(a)** | $0 \text{ min}^{-1}$ |
| $f_M$ | *lag-2* mRNA production rate **in Z1.pp/Z4.aa** | $1.57 \text{ min}^{-1}$ |
| | *lag-2* mRNA production rate **in Z1.ppp/Z4.aaa** | $3.10 \text{ min}^{-1}$ |
| **Model D1** | **Transition from low to high LAG-2 protein degradation** |  |
| $f_D$ | LAG-2 protein production rate **in Z1.pp/Z4.aa** | $0.17 \text{ min}^{-1}$ |
|  | LAG-2 protein production rate in **Z1.ppp/Z4.aaa** | 0.8 min<sup>-1</sup> |
| $b_D$ | LAG-2 protein degradation rate in **Z1.pp/Z4.aa** | 0.017 min<sup>-1</sup> |
|  | LAG-2 protein degradation rate in **Z1.ppp/Z4.aaa** | 0.08 min<sup>-1</sup> |
| **Model D2** | **Transition from low to high LAG-2 protein degradation, non-bursty *lag-2* expression** |  |
| $b_O$ | Transition rate to closed promoter configuration in **Z1.pp(p)/Z4.aa(a)** | 0 min<sup>-1</sup> |
| $f_M$ | *lag-2* mRNA production rate in **Z1.pp/Z4.aa** | 1.57 min<sup>-1</sup> |
|  | *lag-2* mRNA production rate in **Z1.ppp/Z4.aaa** | 3.10 min<sup>-1</sup> |
| $f_D$ | LAG-2 protein production rate in **Z1.pp/Z4.aa** | 0.17 min<sup>-1</sup> |
|  | LAG-2 protein production rate in **Z1.ppp/Z4.aaa** | 0.8 min<sup>-1</sup> |
| $b_D$ | LAG-2 protein degradation rate in **Z1.pp/Z4.aa** | 0.017 min<sup>-1</sup> |
|  | LAG-2 protein degradation rate in **Z1.ppp/Z4.aaa** | 0.08 min<sup>-1</sup> |
| **YFP** | **Dynamics of YFP expression for independent *lag-2* promoter** |  |
| $f_{O_{YH}}$ | Binding of HLH-2 to transgenic *lag-2* promoter | 1·10<sup>3</sup> min<sup>-1</sup> |
| $b_{O_{YH}}$ | Unbinding of HLH-2 from transgenic *lag-2* promoter | 1·10<sup>2</sup> min<sup>-1</sup> |
| $f_{M_Y}$ | *yfp* mRNA production rate in **Z1.pp/Z4.aa** | 3.99 min<sup>-1</sup> |
|  | *yfp* mRNA production rate in **Z1.ppp/Z4.aaa** | 6.32 min<sup>-1</sup> |
| $b_{M_Y}$ | *yfp* mRNA degradation rate | 0.087 min<sup>-1</sup> |
| $f_{D_Y}$ | YFP protein production rate | 0.17 min<sup>-1</sup> |
| $b_{D_Y}$ | YFP protein degradation | 0.017 min<sup>-1</sup> |

## Acknowledgements

We thank Marco Betist (Korswagen lab) for help and advice regarding injections and the integration of the *lag-2* extrachromosomal transgene, Daniel Shaye (Greenwald lab) for advice regarding the *lin-12(0)* line and Kim Renders for help with cloning. Some *C. elegans* strains were provided by the CGC, which is funded by NIH Office of Research Infrastructure Programs (P40 OD010440). NG was partly supported by a fellowship from the Human Frontier Science Program (HFSP) (LT000227/2018-L). The work was supported by a European Research Council Starting Grant (338200-STOCHCELLFATE) awarded to JSvZ.

## Author Contributions

SK, NG and JSvZ conceived the research project; NG, SK, JRK and AK performed time-lapse experiments and data analysis; SK performed smFISH experiments and data analysis; SK and YG performed strain creation; NG and JSvZ performed mathematical modeling; SK, NG and JsvZ wrote the manuscript.

