## Supplementary Figures for "Random birth order and bursty Notch ligand expression drive the stochastic AC/VU cell fate decision in *C. elegans*"

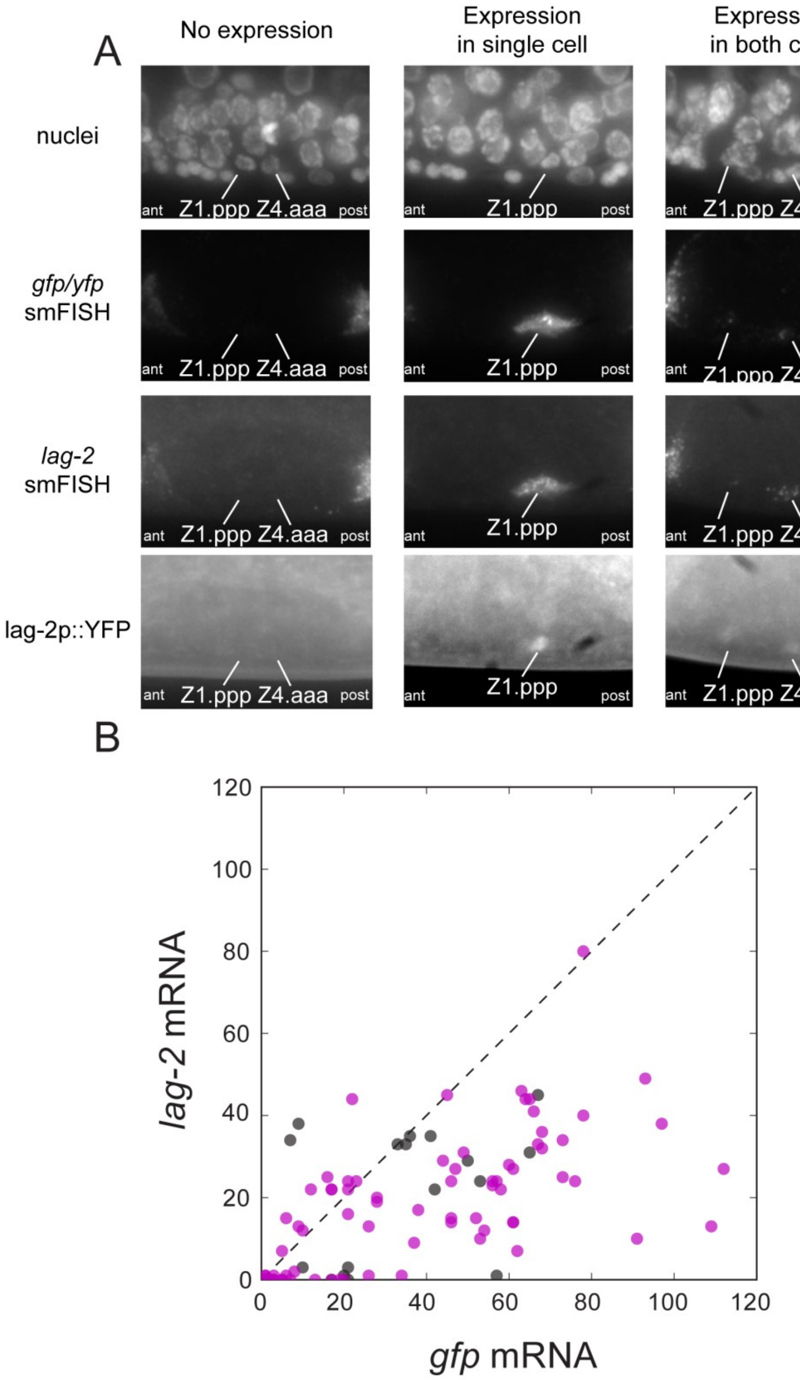


Supplementary Figure 1. **Relationship between *lag-2* and *lag-2p::nls::yfp* expression.**

**(a)** Representative examples of correlated *lag-2p::nls::yfp* and *lag-2 expression* in single animals that show no *lag-2* expression (left column), *lag-2* expression in a single cell (middle column) and *lag-2* expression in both cells (right column). Correlation in expression was examined by simultaneous imaging of *yfp* and *lag-2* mRNA by single molecule FISH and lag-2p::YFP protein levels by YFP fluorescence. *yfp* mRNA molecules were visualized using *gfp* smFISH probes, as they different only by a single point mutation. (b) *yfp* and *lag-2* mRNA levels measured in the same animal. Each marker corresponds to a single animal with color indicating Z1.pp/Z4.aa (black) and Z1.ppp/Z4.aaa cells (magenta).

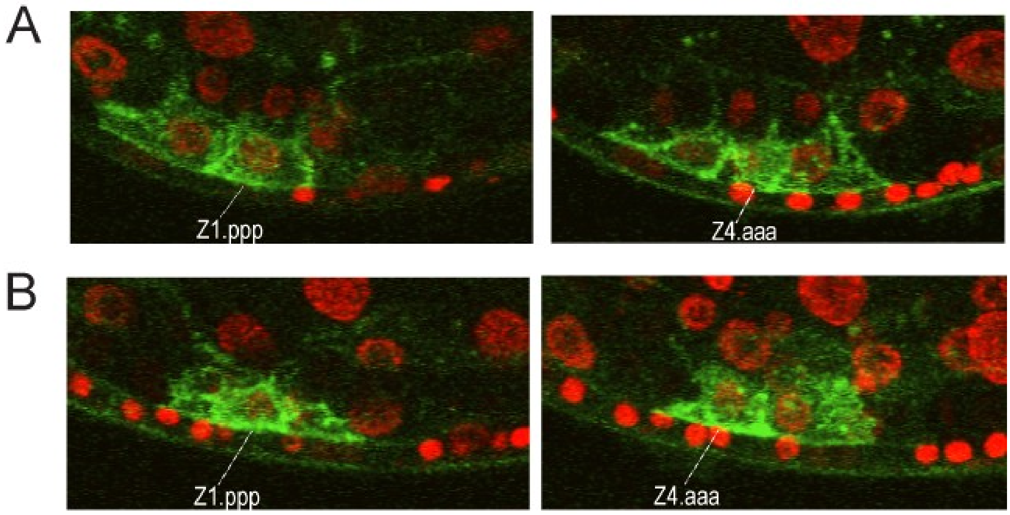


Supplementary Figure 2. **LIN-12::GFP dynamics**

**(a,b)** Representative confocal microscopy images for two *wgIs72[LIN-12::GFP; itIS37[pie-1p::mCherry::H2B::pie-1]]* animals showing LIN-12::GFP (green) and histone (red) fluorescence in Z1.ppp (left column) and Z4.aaa (right column) cells at the late L2 larval stage when the AC/VU decision has been completed. In both animals (a,b), Z1.ppp and Z4.aaa are in different imaging planes and Z4.aaa is identified as the AC by the arrangement of Z1.pp(p/a) and Z4.aa(a/p) cells. Equal LIN-12::GFP fluorescence is observed in Z1.ppp and Z4.aaa cells, indicating that LIN-12 protein levels are not restricted to a single cell.

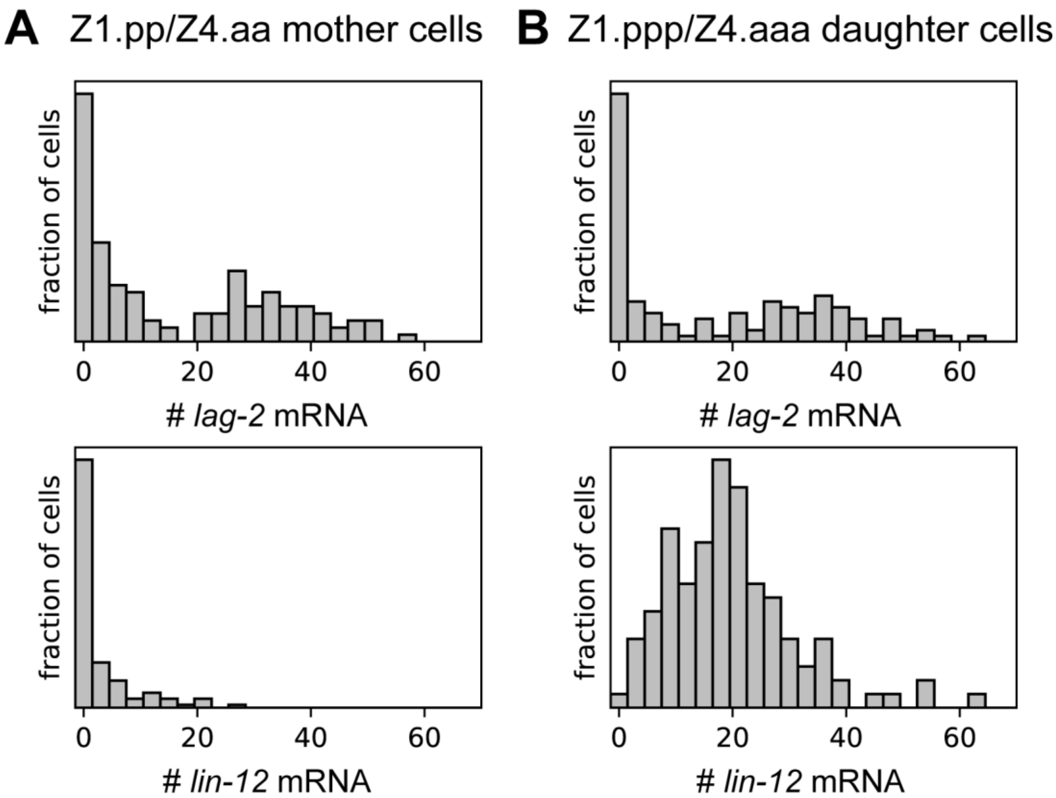


Supplementary Figure 3. **Distribution of *lag-2* and *lin-12* expression in Z1.pp(p) and Z4.aa(a) cells**

**(a,b)** Distribution of *lag-2* (top panel) and *lin-12* (bottom panel) mRNA number obtained by single molecule FISH in (a) mother cells Z1.pp/Z4.aa and (b) daughter cells Z1.ppp/Z4.aaa.

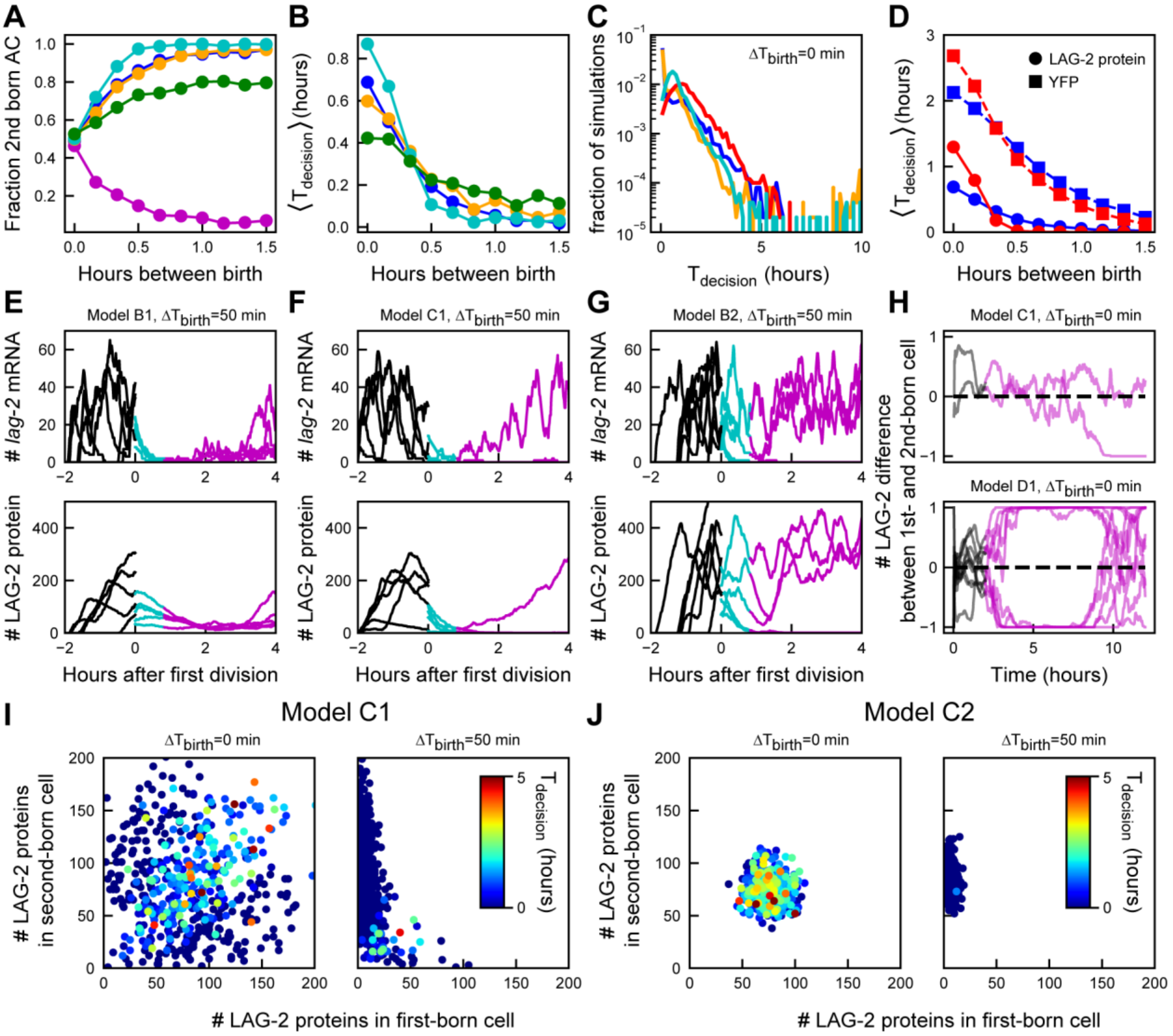


| | Notch activation permitted | LAG-2 protein degradation | *lag-2* expression |
|---|---|---|---|
| Model A1 | *in Z1.pp(p)/Z4.aa(a)* | *constant* | *bursty* |
| Model B1 | *only in Z1.ppp/Z4.aaa* | *constant* | *bursty* |
| Model C1 | *only in Z1.ppp/Z4.aaa* | *Notch-dependent* | *bursty* |
| Model D1 | *only in Z1.ppp/Z4.aaa* | *low in Z1.pp/Z4.aa, high in Z1.ppp/Z4.aaa* | *bursty* |
| Model A2 | *only when both Z1.ppp and Z4.aaa present* | *constant* | *bursty* |
| Model C2 | *only in Z1.ppp/Z4.aaa* | *Notch-dependent* | *non-bursty* |
| Model D2 | *only in Z1.ppp/Z4.aaa* | *low in Z1.pp/Z4.aa, high in Z1.ppp/Z4.aaa* | *non-bursty* |
| Model B2 | *only in Z1.ppp/Z4.aaa* | *constant high* | *bursty* |

Supplementary Figure 4. **Overview of AC/VU models.**

**(a)**, **(b)** Comparing (a) birth order bias and (b) average time to decision $\langle T_{\text{decision}} \rangle$ as a function of the time between Z1.ppp and Z4.aaa birth, $\Delta T_{\text{birth}}$, for the models A2, B2, D1 and D2 (See table for overview of all models). For each $\Delta T_{\text{birth}}$, n=500 simulations. **(c)** Comparing the distribution of decision times for simultaneous Z1.ppp and Z4.aaa birth, for models C1, C2, D1 and D2 ($n$=$5 \cdot 10^3$ simulations). Models D1,2 exhibit longer decision times than models C1,2, particularly for very long, >6 hour, time to decision. **(d)** Average time to decision $\langle T_{\text{decision}} \rangle$ for models C1 (red) and C2 (blue), as determined from LAG-2 protein level (circles) and from YFP expression driven by a second, independent *lag-2* promoter (squares). The lower YFP degradation rate, compared to that of LAG-2, results in longer apparent time to decision, but has minimal impact on the difference in shape of the curve between models C1 and C2. **(e)**, **(f)**, **(g)** Representative trajectories for $\Delta T_{\mathrm{birth}}=50\ \mathrm{min}$ of *lag-2* mRNA (top panel) and LAG-2 protein (bottom panel) dynamics in the first-born cell, for models B1, C1 and B2. For (e) constant, low LAG-2 protein degradation rate (model B1), *lag-2* mRNA is depleted in the first-born cell but LAG-2 protein level remains high, resulting frequently in the first-born cell assuming not VU but rather AC fate, evident by high *lag-2* expression. In contrast, for (f) high LAG-2 degradation rate induced by Notch-signaling (model C1), LAG-2 protein levels are efficiently depleted in the first-born cell. However, for (g) constant, high LAG-2 degradation (model B2), increased fluctuations due to high LAG-2 turn-over result in many first-born cells assuming AC fate. **(h)** Comparing LAG-2 protein dynamics for simulations with exceptionally long, >6 hr, decision times for model C1 (top) and D1 (bottom). Shown is the normalized difference $(D_1\text{-}D_2)/(D_1+D_2)$ where $D_1$ and $D_2$ are the LAG-2 protein numbers in the first- and second-born daughter cell. For model C1, where high LAG-2 protein degradation only occurs in response to Notch signaling, such trajectories represent rare cases where Notch signaling fails to break symmetry between Z1.ppp/Z4.aaa for a long time. In contrast, for model D1, where high LAG-2 degradation occurs in daughter cells independently of Notch signaling, symmetry is initially broken, but spontaneous fluctuations in the *lag-2* expressing AC cause the system to reverse. These fluctuations are caused by the increased turnover of LAG-2 protein in the AC, and do not occur in model C1 as the AC receives no incoming Notch signaling from the neighboring VU cell.

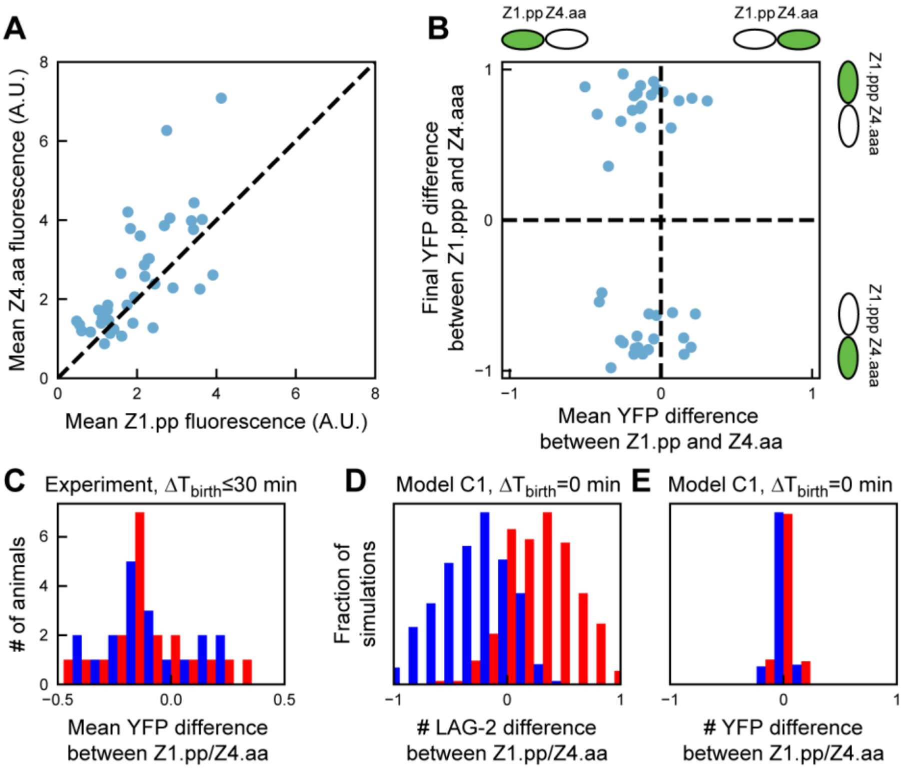


Supplementary Figure 5. **Correlation between Z1.pp/Z4.aa *lag-2* expression and cell fate.**

**(a)** Correlation between YFP expression in mother cells Z1.pp/Z4.aa that divide at similar times ($\Delta T_{\mathrm{birth}}<30$ min). Each marker corresponds to a single animal, with YFP expression in Z1.pp/Z4.aa averaged over 1 hour before cell division. Even though expression levels are weakly correlated in Z1.pp/Z4.aa mother cells, clear stochastic differences in YFP expression occur between both cells. **(b)** Difference in YFP expression in daughter cells Z1.ppp/Z4.aaa at the end of the AC/VU decision, as function of the initial difference in their mother cells Z1.pp/Z4.aa, averaged over 1 hour prior to their division. Only animals are selected that divide at similar times ($\Delta T_{\mathrm{birth}}<30$ min). In absence of a strong difference in birth order, a difference in expression of the *lag-2p::yfp::nls* reporter between mother cells does not provide a bias for the AC/VU decision. For instance, animals in which YFP is expressed more highly in the mother cell Z1.pp are not more likely to have YFP expression, and hence AC fate, restricted to its daughter Z1.ppp. YFP difference is measured as the normalized difference $(Y_1\text{-}Y_4)/(Y_1+Y_4)$, where $Y_1$ and $Y_4$ are the YFP fluorescence intensity in Z1.pp(p) and Z4.aa(a) respectively. **(c)** Histogram of the normalized difference in YFP expression between mother cells Z1.pp/Z4.aa, average 1 hour prior to division, for animals where Z1.ppp (red) or Z4.aaa (blue) assume AC fate. Both distributions overlap fully, indicating that relative YFP level in mother cells does not correlate with cell fate outcome. **(d), (e)** Histogram of the normalized difference in (d) LAG-2 protein and (e) YFP between mother cells Z1.pp/Z4.aa just prior to division, in simulations of model C1 where mother cells divide simultaneously. Color indicates whether Z1.ppp (red) or Z4.aaa (blue) assumed AC fate. In agreement with the experimental data in (c), no correlation between YFP difference and cell fate outcome is observed. However, difference in LAG-2 protein levels between Z1.pp/Z4.aa do correlated with outcome, with the mother cell showing the highest LAG-2 expression more likely to assume AC fate. This is because bursty *lag-2* and *yfp* expression are uncorrelated, meaning that a mother cell with high YFP levels could have low LAG-2 protein level.

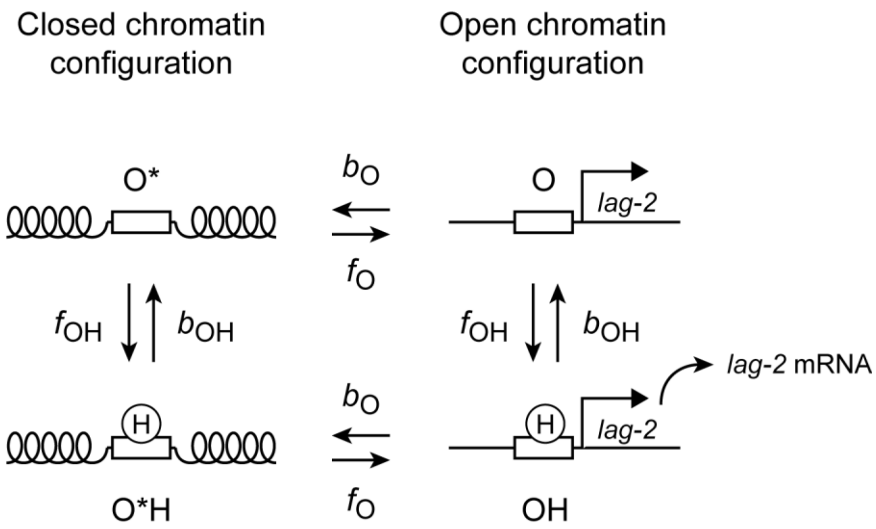


Supplementary Figure 6. **Model of the *lag-2* promoter.**

The *lag-2* promoter is assumed to exist in an open configuration $O$ and closed configuration $O^*$, with transitions from the closed to open configuration with forward rate $f_O$ and backward rate $b_O$. HLH-2, the transcriptional activator of lag-2, can bind the promoter both in the closed and open configuration with forward rate $f_{OH}$ and backward rate $b_{OH}$. Moreover, transitions between the open and closed configuration occur independent of whether HLH-2 is bound. However, transcription of lag-2 mRNA only occurs from the open promoter configuration with HLH-2 is bound to it. For sufficiently slow transitions between open and closed configurations, this model gives rise to bursty *lag-2* expression, but only in the presence of HLH-2.
